# Excitatory Neuron-Derived Interleukin-34 Controls Cortical Developmental Microglia Function

**DOI:** 10.1101/2024.05.10.589920

**Authors:** Benjamin A. Devlin, Dang M. Nguyen, Diogo Ribeiro, Gabriel Grullon, Madeline J. Clark, Amelie Finn, Alexis M. Ceasrine, Seneca Oxendine, Martha Deja, Ashka Shah, Shomik Ati, Anne Schaefer, Staci D. Bilbo

## Abstract

Neuron-microglia interactions dictate the development of neuronal circuits in the brain. However, the factors that regulate these processes across development are largely unknown. Here, we find that IL34, a neuron-derived cytokine, is upregulated in early development and maintains neuroprotective, mature microglia in the anterior cingulate cortex (ACC) of mice. We show that IL34 is upregulated in the second week of postnatal life and is expressed primarily in excitatory neurons. Excitatory-neuron specific knock-out of IL34 reduced microglia number and TMEM119 expression and increased aberrant microglial phagocytosis of excitatory thalamocortical synapses in the ACC. Acute, low dose blocking of IL34 at postnatal day 15 similarly decreased TMEM119 and inappropriately increased microglial phagocytosis of synapses. Viral overexpression of IL34 induced TMEM119 expression and prevented appropriate microglial phagocytosis of synapses. These findings establish IL34 as a key regulator of neuron-microglia crosstalk in postnatal brain development, controlling both microglial maturation and synapse engulfment.

## Introduction

Microglia serve diverse roles in normal brain development including functions in vasculogenesis, neurogenesis, myelination, tissue repair, and synaptic remodeling^1^. Microglia are tissue-resident macrophages and progress through different functional stages in development^2^ dependent upon precise molecular cues they receive from other cells in their neural environment. These cues are still being defined and likely depend on the distinct brain region and/or specific circuits within a given region. In many brain regions, it is not until the second postnatal week of life in mice that microglia adopt a transcriptional and functional profile consistent with adult microglia^2,3^; however, little is known about the precise timing of this transition, or the brain-specific signals that contribute to it^4^.

As macrophages, microglia depend on signaling through the colony-stimulating factor 1 receptor (CSF1r) for their proliferation, differentiation, and survival^5–7^. For this reason, CSF1r is often targeted for microglial depletion experiments^8^, and blockage or genetic knock-out of this receptor potently reduces microglia number in the brain at every stage of development^9^. Often overlooked is that there are two known ligands that signal through this receptor: CSF1, which is expressed in all tissues, and the more recently identified interleukin-34 (IL34), whose expression is limited to keratinocytes in the skin and neurons in the forebrain^10,11^. Further, there is a spatio-temporal pattern of microglial dependence on, and presence of, CSF1 and IL34. In the embryonic and early postnatal mouse brain, CSF1 is predominantly expressed in all brain regions, and the loss of CSF1 alone is sufficient to deplete virtually all microglia. This is in contrast to the adult brain, where IL34 is expressed at higher levels in the forebrain; thus roughly 70-90% of gray matter microglia in the forebrain depend on IL34 signaling for survival, while microglia in the cerebellum and brainstem still depend on CSF1^7,10–13^. It is notable that these signals come from different cellular sources, as CSF1 is primarily expressed by microglia early in life^2^ and later by oligodendrocytes and astrocytes, while IL34 is expressed by neurons^13^. Thus, given these differences in timing, brain region, and cell source, it is highly unlikely these two ligands play a redundant role for microglial survival and differentiation, however no work to date has empirically tested this *in vivo*. There does exist some preliminary evidence that CSF1 and IL34 drive distinct transcriptional programs in microglia^14^, but it is still unknown (1) when forebrain microglia become primarily dependent on IL34, (2) how neurons, and which neurons, express IL34, and (3) how IL34 could differentially influence microglia cell state and function, *in vivo*, in cortical development.

Here, we demonstrate that IL34 is developmentally upregulated in the second week of postnatal life in mice in multiple forebrain regions. We show that IL34 expression in the cortex is regulated by neuronal subtype, with a higher level in glutamatergic (VGlut1+) neurons compared to GABAergic (Gad2+) neurons. Constitutive genetic knock-out of IL34 impacts microglia numbers and developmental cell state as demonstrated by decreased TMEM119 protein and increased lysosomal content at P15. Excitatory-specific IL34 knock-out in VGlut2^Cre^;IL34^fl/fl^ mice impacts microglia similarly to global knock-out and causes increased phagocytosis of VGlut2+ synaptic material in the ACC. Acute, low-dose antibody blocking of IL34 at P15 prevents drastic microglia loss but decreases the amount of homeostatic (TMEM^Hi^/CD68^Lo^) microglia, increases phagocytic microglia (TMEM^Lo^/CD68^Hi^), and increases aberrant microglial phagocytosis of synapses. Finally, viral overexpression of IL34 in neurons at a developmentally inappropriate timepoint (P1-P8) prematurely increases microglial TMEM119 and decreases microglial engulfment of synapses.

## Results

### IL34 expression is primarily regulated by development and neuronal subtype

We examined the time course of IL34 and CSF1 expression in the developing mouse brain. Previous literature demonstrated that microglia in the embryonic brain depend solely on CSF1 signaling for survival, whereas microglia in the adult brain depend on either IL34 or CSF1, depending on brain region^7,10–12^. Despite this, little was known about when IL34 becomes the primary signal for microglia in regions that show high levels of adult IL34 expression (e.g. cortex and striatum)^13^. We collected brains from wild-type, C57Bl6/j male and female mice at 6 ages (postnatal day 7, 14, 20, 30, 38, 55) spanning early life, adolescence, and early adulthood in mice^15^. We measured IL34 mRNA in three IL34-dependent brain regions (Anterior Cingulate Cortex (ACC), Nucleus Accumbens (NAc), Amygdala (AMY)) and from the cerebellum (CBM), which is CSF1-dependent (Figure 1A). We found no developmentally regulated expression of *CSF1* mRNA in any region, but a striking developmental increase in *IL34* mRNA, which began between postnatal day 7 (P7) and P14 in all regions except the cerebellum (Figure 1B). We collected mice at P8 and P15 and used ELISA to measure total IL34 protein in the ACC and CBM and similarly found an increase in protein between P8 and P15 in the ACC, but no increase in the cerebellum (Figure 1C). We focus on ACC in all following experiments, as previous work in our lab has detailed a developmental period of microglia-neuron interactions in this brain region^16^. Together, IL34 expression increases in IL34-dependent brain regions in the second postnatal week.

**Figure 1.**
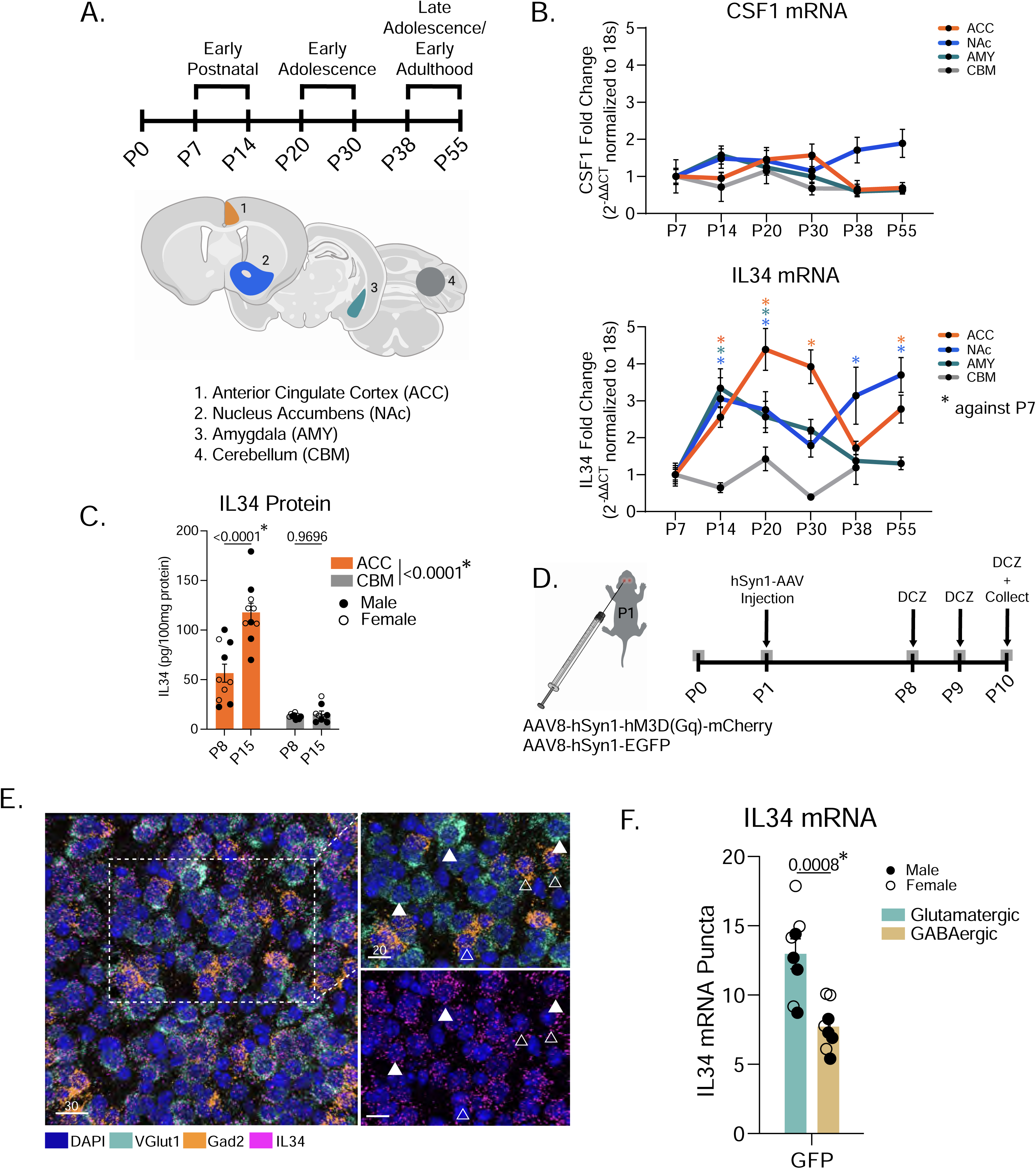
IL34 expression is primarily regulated by development and neuronal subtype. (A) Experimental timeline for WT brain collection across development and brain regions punched for qPCR analysis. (B) Quantification of IL34 and CSF1 mRNA levels in all four brain regions across six postnatal developmental timepoints. (n = 8 male and 8 female C57Bl6/j wild-type mice/age, P55 cerebellum excluded due to poor RNA quality, one-way ANOVA, data normalized to P7). (C) Quantification of IL34 protein using ELISA from tissue punches of the ACC and CBM. (n = 5 male and 5 female C57Bl6/j wild-type mice/age, two-way ANOVA age x brain region, Sidak’s *post hoc* test, main effect of brain region in legend). (D) Experimental timeline for early postnatal viral injections and DREADDs experiments. (E) Representative image of RNA-FISH stain including VGlut1 to distinguish excitatory neurons from inhibitory neurons. (F) Quantification of IL34 mRNA puncta in excitatory (VGlut1+) and inhibitory (Gad2+) neurons in control mice. (n = 4 mice/sex, data shown are animal averages of the expression level of all neurons of that type, unpaired t-test). Related to Figure S1.

To test whether neuronal activity or subtype influences IL34 expression, we artificially increased neuronal activity using a designer receptor exclusively activated by designer drug (DREADDs) approach in the first postnatal week (Figure 1D). We injected the designer drug deschloroclozapine (DCZ) at postnatal day 8, 9, and 10. This approach reliably activated neurons in Gq-expressing mice compared to GFP controls as measured by increased Fos expression (Figures S1A-S1D). Using RNAScope, we stained and quantified IL34 mRNA and found no difference in total IL34 levels between GFP control and Gq mice, suggesting that heightened neuronal activity does not increase overall levels of IL34 in the circuit (Figures S1E-F). We quantified IL34 at single-cell resolution using mRNA probes for *Fos* and *Gad2* to distinguish recently active (*Fos*+, i.e. greater than 6 *Fos* puncta per cell adjusted for cell size) from inactive (*Fos*-), GABAergic (*Gad2*+) or glutamatergic (*Gad2*-) neurons (Figure S1G). We found higher expression of *IL34* in *Fos*+ (active) neurons in Gq, but not GFP, mice. (Figure S1H). Notably, we observed twice as much *IL34* in *Gad2*- (presumably excitatory) neurons versus *Gad2*+ inhibitory neurons. Thus, to follow-up, we used *VGlut1* and *Gad2* probes to precisely quantify *IL34* in glutamatergic (*VGlut1*+) and GABAergic neurons and confirmed that *IL34* is expressed two-fold higher in excitatory neurons compared to inhibitory neurons (Figures1E-F). We next tested whether activating only excitatory neurons impacted *IL34* expression using VGlut1-Cre mice and a cre-dependent DREADD virus (Figure S1I). We again saw no change in overall *IL34* (Figures S1J-K), but an increase in *IL34* in *Fos*+ neurons compared to *Fos*- neurons (Figure S1L). These data suggest there is a homeostatic setpoint for *IL34* that is unchanged by activation, while at the level of individual neurons, activity predisposes them to increase *IL34*. It is worth noting that we did not measure protein levels or neuronal release, which may be impacted by activation and occlude our findings within cell somas.

### Constitutive IL34 knock-out (KO) decreases adult microglial number and TMEM119 protein

Previous reports show that adult mice lacking IL34 (IL34^LacZ/LacZ^) have fewer microglia in the cortex, striatum, and hippocampus, but not the cerebellum or brainstem^10,11^. We confirmed these findings in IL34^LacZ/LacZ^ adult male and female mice, and observed a moderate, yet significant, reduction in Iba1+ cell number in IL34^LacZ/+^ heterozygous mice (Figure S2A-B). Microglial numbers in the cortex and striatum of IL34 KO mice were equivalent to cerebellum and brainstem numbers, suggesting that, when IL34 is lost, a CSF1-dependent population of microglia is maintained in these regions^13^. Interestingly, we found a reduction in TMEM119, a microglia-specific marker that is developmentally regulated and present in mature, homeostatic microglia^3,17^ in the remaining microglia of IL34^LacZ/LacZ^ mice, suggesting that these forebrain, CSF1-dependent microglia are immature and non-homeostatic (Figure S2C). This diminished TMEM119 in IL34 KO microglia was similar to the low expression seen in CSF1-maintained brainstem and cerebellar microglia (Figure S2D). In line with the hypothesis that there is a developmental consequence of IL34 loss on forebrain microglia, we re-analyzed bulk RNA-Sequencing on isolated microglia from WT and IL34 KO mice^14^ and found IL34 KO microglia have elevated levels of neonatal marker genes (*ITGAX, APOE, CLEC7A*) and comparably low expression of adult markers (*P2YR12, TMEM119, OLFML3*)^2,3,18^ (Figure S2E). Notably, IL34 KO microglia have an upregulation of CSF1, which highly expressed in immature microglia^2,19^, and suggests that, in the absence of the primary signal (IL34), this CSF1 upregulation is a potential compensatory survival mechanism. These gene expression changes were not observed in CSF1 KO mice^14^. Additionally, many genes associated with microglia phagocytosis were dysregulated in forebrain IL34 KO microglia (*C3AR1, ITGAM, AXL*)^20–22^(Figure S2E. These data suggest that when IL34 signaling is lost, remaining CSF1-dependent microglia in the forebrain resemble cerebellar microglia and have potentially disrupted phagocytic capabilities. This supports the idea that there are distinct IL34- and CSF1-maintained subpopulations of microglia that are transcriptionally and functionally distinct.

#### Global IL34 KO impacts microglia state in the second postnatal week

Our observation that IL34 increases in the second postnatal week coincides with many known developmental microglial processes, including proliferation^16^, upregulation of TMEM119^3^, increased ramification^23^, establishment of region-specific heterogeneity^24^, and closure of a period of synaptic pruning in the ACC^16^. To test whether IL34 functionally controls these processes, we generated and collected IL34^LacZ/+^ (heterozygous control) and IL34^LacZ/LacZ^ (KO) mice at P8 and P15 (Figure S3A). We confirmed the developmental increase in IL34 by staining for LacZ in IL34^LacZ/+^ mice and saw robust expression at P15 (Figure 2A). We next stained the ACC with Iba1 and TMEM119 and found that IL34 KO mice have a reduction in microglia at P8 and P15 and a reduction in TMEM119 protein relative to Iba1 in IL34 KO microglia at P15 (Figure 2B-E), suggesting that IL34 is necessary for the developmental upregulation of TMEM119 in cortical microglia. We also observed a marked reduction in expression of another homeostatic protein (P2YR12) at both P8 and P15 (Figure S3B). To assess whether IL34 KO microglia have dysregulated phagocytic function we measured the volume of CD68 (lysosomal marker) within microglia (Iba1) and found that P15, but not P8, microglia in IL34 KO mice have elevated lysosomal content (Figure 2F-G). Cell volume was unchanged in IL34 KO mice (Figure S3C). We also found a decrease in process complexity in IL34 KO microglia at P15 (Figure 2H-J). These results suggest that IL34’s role in the brain extends beyond survival and has developmentally relevant functions for microglial proliferation, maturation, phagocytosis, and ramification.

**Figure 2.**
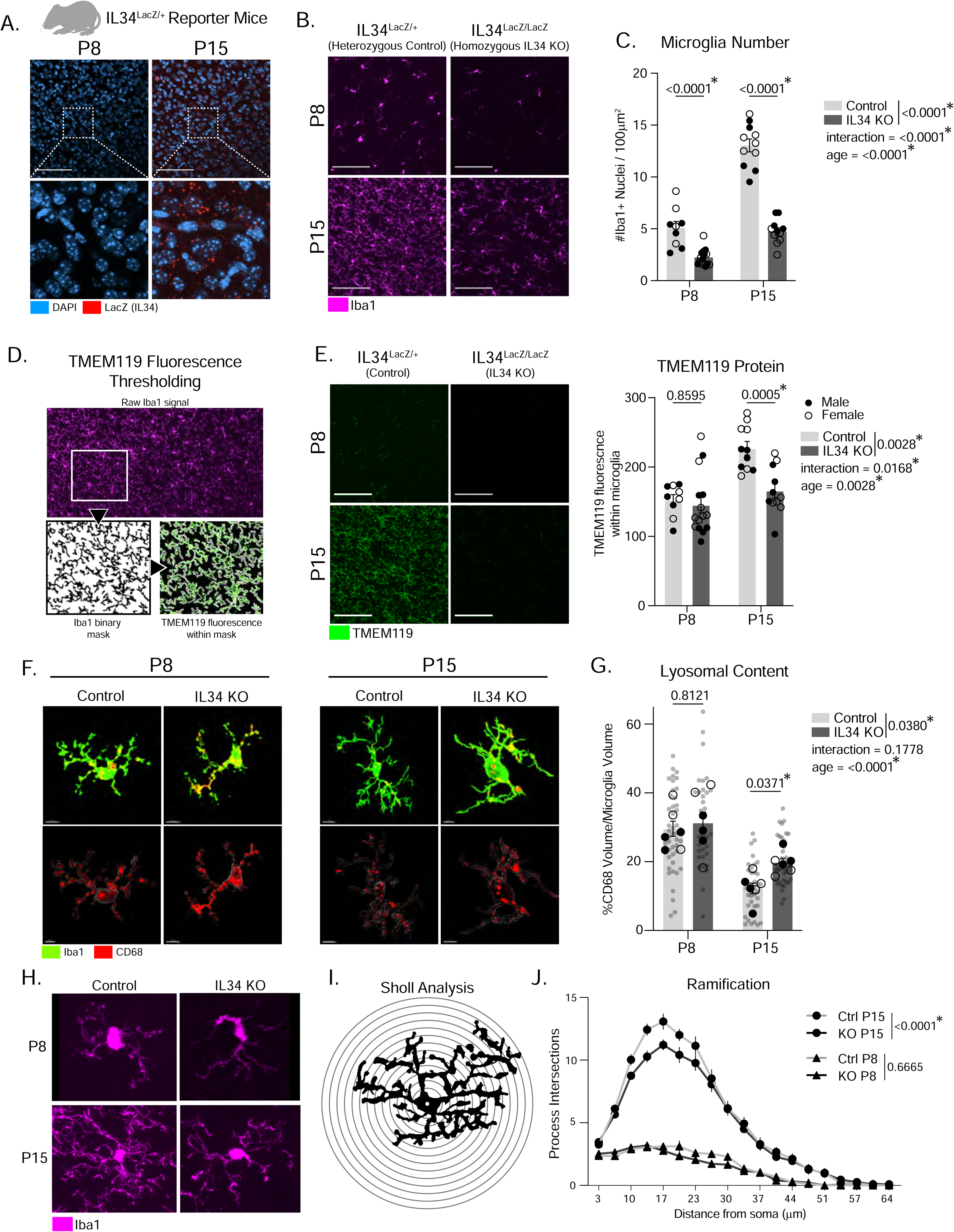
Constitutive IL34 KO impacts microglia in the second postnatal week. (A) Representative image of LacZ in the ACC of IL34^LacZ/+^ mice at postnatal day 8 (P8) and P15. (B-C) Representative images of Iba1 and quantification of microglia number in the ACC of P8 and P15 IL34^LacZ/LacZ^ and IL34^LacZ/+^ mice. (n = 5-8 mice/sex/age/genotype, two-way ANOVA age x genotype, Sidak’s *post-hoc* test, main effect of genotype and interaction in legend). Scale = 100µm. (D) Method for masking the Iba1 channel to quantify TMEM119 fluorescence. (E) Representative images and quantification of TMEM119 in the ACC of P8 and P15 IL34 KO and control mice (n = 5-8 mice/sex/age/genotype, two-way ANOVA age x genotype, Sidak’s *post-hoc* test, main effect of genotype and interaction in legend). (F) Representative IMARIS 3D reconstructions of microglia (Iba1) and lysosomes (CD68) from ACC of P8 and P15 IL34 KO and control mice. Scale = 5µm for P8 microglia, 15µm for P15 microglia. (G) Quantification of lysosomal content (volume of CD68 / total microglia volume * 100, n = 3 mice/sex/age/genotype, 4-6 cells analyzed per mouse, individual microglia represented by gray circles, animal averages represented by black dots, two-way ANOVA age x genotype, Sidak’s *post-hoc* test, main effect of genotype and interaction in legend). (H-J) Representative images of individual microglia (Iba1) and quantification of ramification using sholl analysis. (n = 5-8 mice/sex/age/genotype, 6 cells analyzed per mouse, two-way ANOVA distance from soma x genotype, main effect of genotype in legend). Related to Figure S2-S4.

We found no effect of IL34 KO on microglia number, TMEM119 protein, or P2YR12 protein in the brainstem, a CSF1-dependent region (Figure S3D-F). We also found no change in neuron, astrocyte, or oligodendrocyte cell numbers (Figure S3H-J) in ACC, and bulk RNASequencing of whole forebrain from IL34 KO and control mice demonstrated downregulation of microglia genes (e.g. *P2YR12, CSF1R, TMEM119*) but no changes in non-microglial genes (Figure S3K). These data confirm that IL34 genetic KO impacts microglia in the forebrain and not the brainstem, and does not affect the number or transcriptional profile of neurons, astrocytes, or oligodendrocytes^5,10^.

#### Global IL34 KO imparts an anxiolytic phenotype at P15 and adulthood

To test whether the functional changes to microglia influenced behavioral outcomes (Figure S4A), we first tested neonatal ultrasonic vocalizations (USVs), which have been shown to be developmentally regulated in the second week of postnatal life and impacted by early-life microglial perturbations ^16,25^. We found no difference in call number at P8 but a decrease in P15 KO mice (Figures S4B-C). IL34 KO mice also have increased weight at P8 and P15 (Figures S4F-G) and they spent an increased percentage of time in the open arms of the elevated plus maze (EPM) in adulthood (Figures S4D-E), suggesting an anxiolytic phenotype. There was no difference in locomotor activity or sociability (Figure S4H-I), but IL34 KO males made more errors during Barnes Maze reversal (Figure S4J-K), suggesting they have a moderate deficit in cognitive flexibility. Taken together, IL34 KO induces an anxiolytic phenotype at P15 (reduced vocalizations) that persists to adulthood (increased open arm time in the EPM). These findings make sense given the ACC’s role in regulating anxiety^26^, and suggest that the microglia changes in IL34 KO mice impact behavior early in life and in adulthood.

#### Excitatory neuron-specific IL34 KO decreases microglia number and increases phagocytosis of synapses

To functionally link excitatory neuron-derived IL34 to the microglia changes we observed in the global KO mice, we generated VGlut2^Cre^;IL34^fl/fl^ mice and collected their brains at P15. This VGlut2^Cre^ genetic strategy predominantly affects excitatory thalamic neurons, thus testing whether IL34 released from thalamic projections, i.e. synaptic terminals, into the ACC was sufficient to influence cortical microglia maturation and function. Comparable to the global IL34 KOs, we found a reduction in microglia number and TMEM119 protein in VGlut2^Cre^;IL34^fl/fl^ mice (Figure 3A-C).

**Figure 3.**
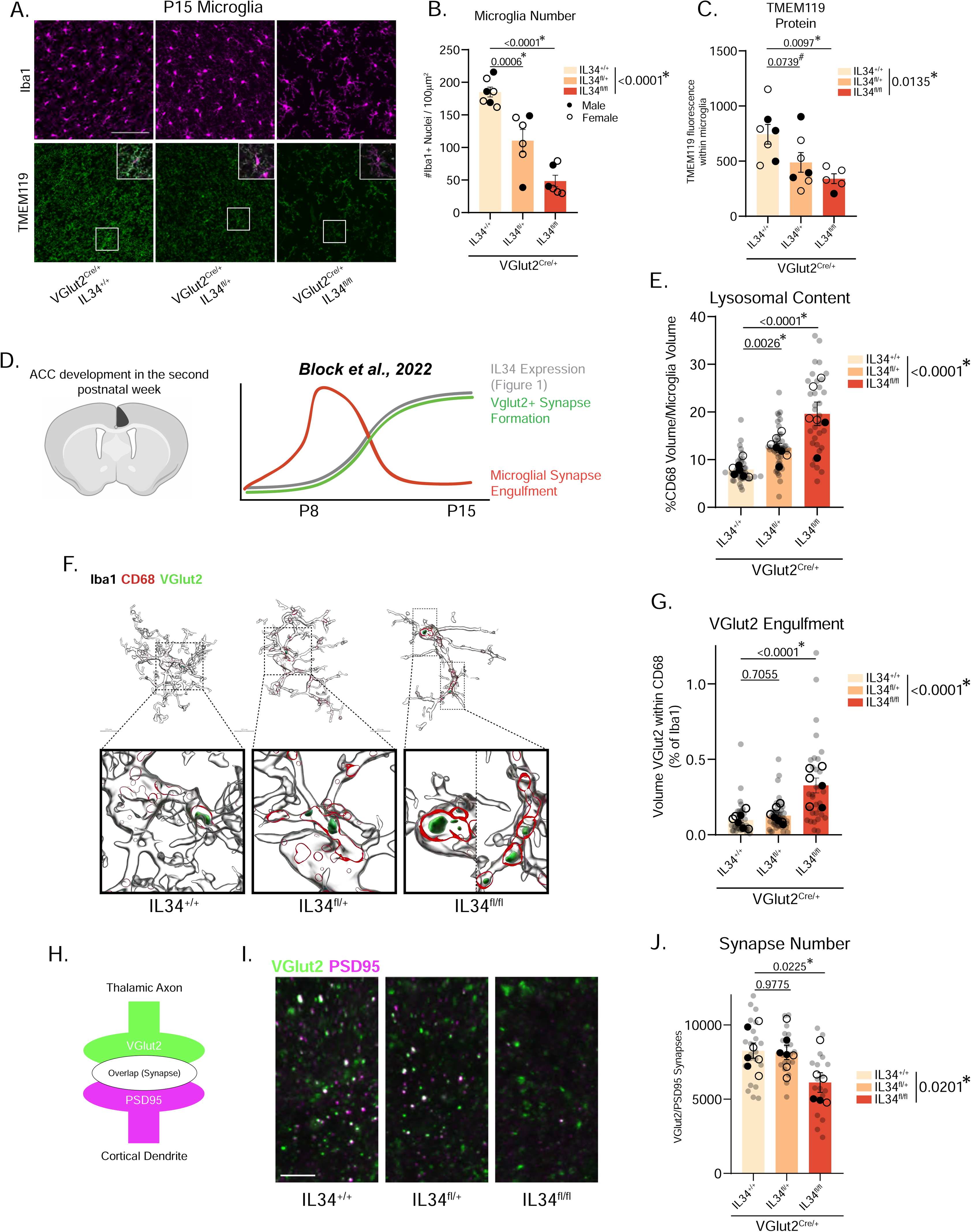
Excitatory neuron-specific KO of IL34 reduces microglia number, increases phagocytosis of synaptic material, and reduces overall synapse numbers in the ACC at P15. (A) Representative images of Iba1 and TMEM119 in the ACC of P15 VGlut2^Cre^ IL34^+/+^, IL34^fl/+^, and IL34^fl/fl^ mice. Scale = 100µm. (B-C) Quantification of microglia number and TMEM119 in the three groups. (n = 2-4 mice/sex/genotype, one-way ANOVA, Sidak’s *post-hoc* test, main effect of genotype in legend). (D) Previous findings detailing microglia-neuron interactions in the ACC during the second postnatal week. Conceptual diagram constructed based on data from Figure 4 in Block et al., 2022. (E) Quantification of total lysosomal content in the ACC of P15 VGlut2^Cre^ IL34^+/+^, IL34^fl/+^, and IL34^fl/fl^ mice. (n = 2-4 mice/sex/genotype, 4-6 cells analyzed per mouse, individual microglia represented by gray circles, animal averages represented by black dots, nested one-way ANOVA, Sidak’s *post-hoc* test, main effect of genotype in legend). (F) Representative IMARIS reconstructions and quantification of microglia synaptic engulfment from VGlut2Cre IL34+/+, IL34fl/+, and IL34fl/fl mice. (n = 2-4 mice/sex/genotype, 4-6 cells analyzed per mouse, individual microglia represented by gray circles, animal averages represented by black dots, nested one-way ANOVA, Sidak’s *post-hoc* test, main effect of genotype in legend). (G) Thalamocortical synapse numbers are quantified by VGlut2 and PSD95 overlap. (I-J) Representative images and quantification of Vglut2+/PSD95+ overlap. (n = 2-4 mice/sex/genotype, 3 sections imaged per animal, images represented by gray circles, animal averages represented by black dots, nested one-way ANOVA, Sidak’s *post-hoc* test, main effect of genotype in legend). Scale = 5µm.

It is well established that microglia engulf aberrant or unnecessary synapses in neuronal development. This process requires precise regulation to ensure appropriate synapses are eaten, while necessary synapses are maintained^27^. While some signals have been identified that act as “eat me” tags on synapses destined for destruction^22,28–31^, very few “don’t eat me” signals have been discovered that inhibit synaptic engulfment, protecting structurally mature, functioning circuits^32^. Recent work from our lab has shown that microglia in the ACC at P8 exhibit elevated phagocytic capacity, actively engulf VGlut2+ thalamocortical presynaptic material, and stop by P10^16^ (Figure 3D). Because this timing coincides with the IL34 increase (P8-P15, Figure1), and because decreasing IL34 induces phagocytic, immature microglia (Figure 2F-G), we hypothesized that IL34 may act as a “brake” on microglial synaptic pruning. To quantify engulfment of thalamocortical synapses (which are the neurons affected in our VGlut2^Cre^;IL34^fl/fl^ KO strategy), we triple-stained for Iba1, CD68, and VGlut2 and 3D reconstructed individual microglia in IMARIS. We found elevated lysosomal content in IL34^fl/fl^ mice (Figure 3E), and increased engulfment of excitatory presynaptic material in IL34^fl/fl^ mice compared to IL34^fl/+^ and IL34^+/+^, confirming our hypothesis that neuronal IL34 prevents microglial engulfment of synapses (Figures3F-G and S4L). To check if microglial over-engulfment led to less excitatory synapses in the ACC, we stained for presynaptic VGlut2 and postsynaptic PSD95 and counted overlap of the two signals (Figure 3H). We found the expected decrease in IL34^fl/fl^ mice (Figure 3I-J), providing strong initial evidence that excitatory neuron-derived IL34 acts as a “don’t eat me” signal for thalamocortical synapses in the ACC.

#### Acute IL34 inhibition at P15 mimics constitutive genetic loss of IL34 and increases phagocytic microglia

One limitation of our findings thus far is that there is a significant reduction in the number of microglia in IL34 KO mice (Figures2B-C and 3C). Thus, it is reasonable to assume that the remaining microglia increase their phagocytic activity to compensate for the fact that there are less of them. To control for this and any off-target developmental effects of the KOs, we injected function blocking antibodies directly into the brain intracerebroventricularly (ICV) to block IL34 (anti-IL34), CSF1 (anti-CSF1), or control IgG^12^ in a temporally precise manner at P15 (Figure 4A). We titrated antibody dose (1mg/kg in our study vs. 100mg/kg in original publication^12^) to reduce microglial cell death. We collected these mice 48 hours after surgery and found only a slight, non-significant decrease in microglia number in the ACC (Figure 4B-C), while there was still a significant decrease in TMEM119 in anti-IL34-treated mice (Figure 4D). The anti-IL34 microglia were also less ramified (Figures4E-F).

**Figure 4.**
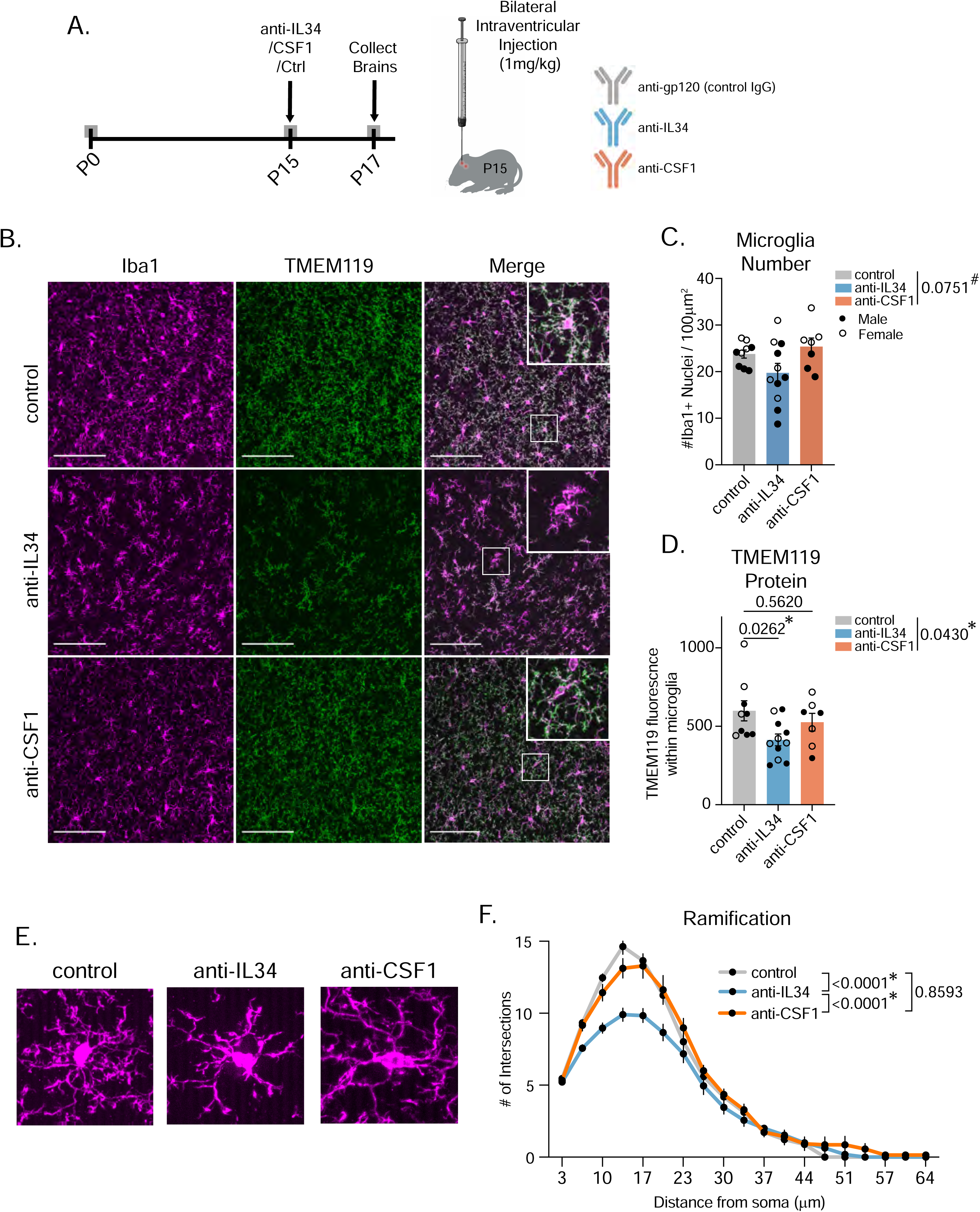
Acute, low-dose IL34 inhibition at P15 mimics constitutive genetic loss of IL34 without significant cell loss. (A) Schematic of blocking antibody injection surgeries and tissue collection. (B) Representative images of Iba1 and TMEM119 in the ACC of mice administered either control (IgG) antibody, anti-IL34, or anti-CSF1. Scale = 100µm. (C-D) Quantification of microglia numbers and TMEM119 in the three groups. (n = 3-6 mice/sex/antibody, one-way ANOVA, Sidak’s *post-hoc* test, main effect of antibody shown in legend). (E) Representative images of individual microglia (Iba1) and quantification of cell ramification from control, anti-IL34, and anti-CSF1 mice. (n = 5-8 mice/sex/treatment, 6 cells analyzed per mouse, two-way ANOVA distance from soma x genotype, main effect of genotype in legend). Related to Figure S7.

We next tested whether the decreased TMEM119 in anti-IL34 microglia correlated with increased lysosomal activity. We noticed significant heterogeneity where some microglia were abnormally high in CD68 (suggestive of heightened phagocytic activity) and low in TMEM119 (Figure 5A). We quantified this by measuring expression of the two markers on a per cell basis and binning them into three categories: “Lo”, “Mid”, and “Hi” based on fluorescent value histograms (Figures S5A-B). Anti- IL34 mice have fewer CD68^Lo^TMEM119^Hi^ microglia, an expression profile typical of homeostatic cortical microglia at P15, and more CD68^Hi^TMEM119^Lo^ phagocytic, non-homeostatic microglia (Figures5B-C). There was also a decrease in TMEM119^Hi^ cells but no change in CD68^Hi^ cells (Figures S5C-D). We then sequenced isolated microglia and whole forebrain from control and anti- IL34 mice (Figure 5D); anti-IL34 treated microglia transcriptomes clustered separately from control microglia via principal component analysis (Figure 5E) and there were over 400 differentially expressed genes (DEGs) between these two groups. Several mature, homeostatic marker genes (e.g. *P2YR12, TMEM119, CD164*) were upregulated in control microglia while anti-IL34 microglia adopted an interferon-responsive inflammatory profile (*CXCL99, IFIT3B, CCL5, MX1*). We observed moderate enrichment for “disease-associated” microglia genes in a-IL34 microglia^33^ (Figures S5E-F), but the gene signature was most similar to type-I-interferon-responsive microglia that shape cortical development in the first week of postnatal life^34^ (Figures5F and S5G). We calculated the “microglia developmental index” using our previously published methods^19^ and found that anti-IL34 treated microglia have reduced transcriptional maturity (Figure 5G). Comparing whole forebrain differences revealed a similar DEG pattern to the isolated microglia, confirming that IL34-blocking induces changes specific to microglia, and not astrocytes, oligodendrocytes, or neurons, all of which have been shown to express receptors for IL34 at low expression and affinity levels^35–38^ (Figure 5H). Anti- IL34 whole forebrain samples also have enrichment for genes related to microglial response to Lipopolysaccharide (LPS), which make up an “Inflammation Index”^19^ (Figure 5I). This implies reduced IL34 contributes to brain-wide inflammation. Together, preventing IL34 signaling acutely at P15 phenocopies the effects of genetic IL34 KO, inducing phagocytic, immature microglia. We acknowledge the limitations that come with injecting blocking antibodies, which causes tissue damage and microglia activation. However, we feel confident in these data because they corroborate findings from the non-invasive, genetic IL34 KO experiments, and we observed no tissue damage in or around our analysis region as the injections were 2mm posterior to the ACC (Figure S6A).

**Figure 5.**
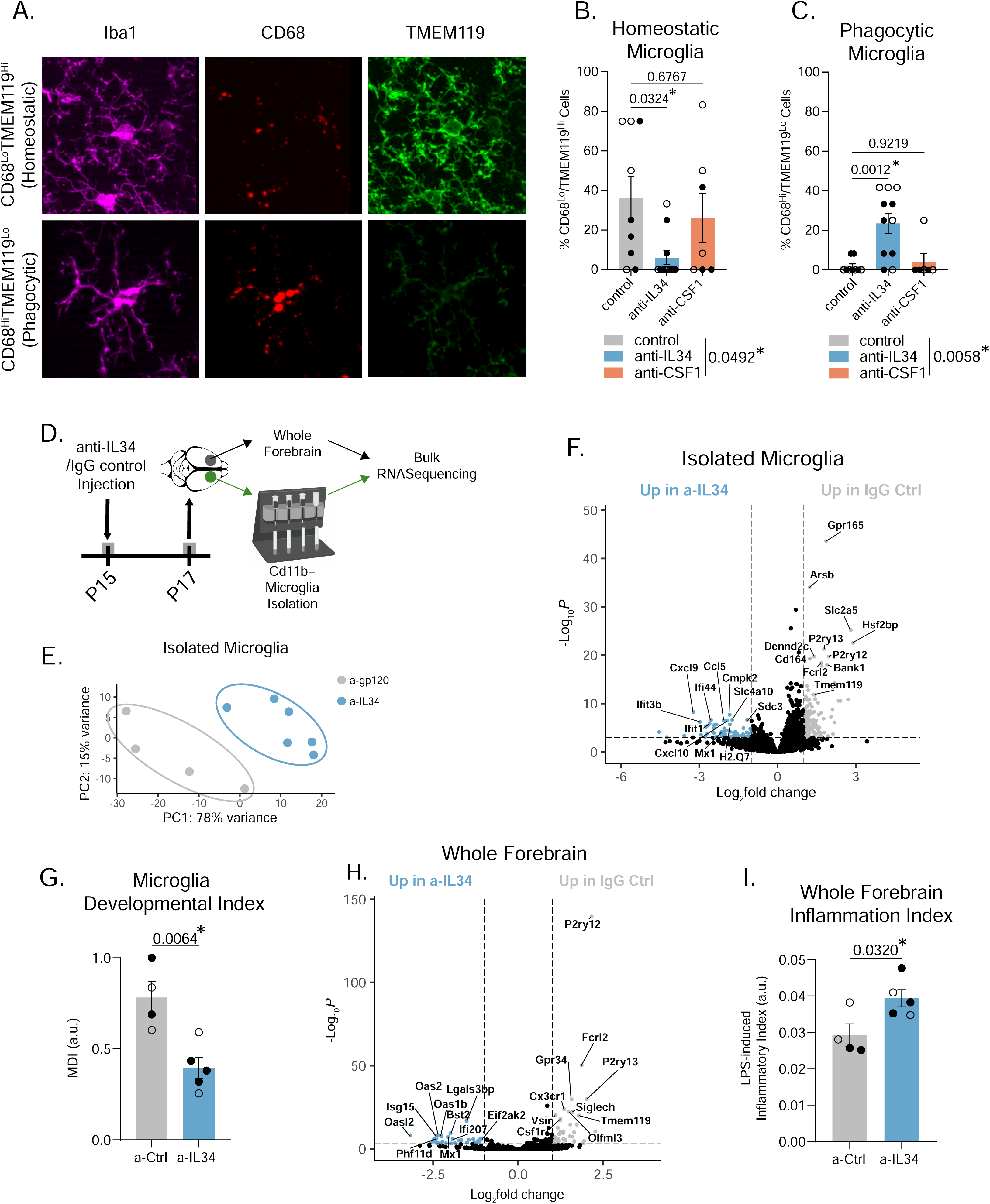
Acute IL34 inhibition at P15 decreases mature, homeostatic microglia and increases phagocytic microglia. (A) Representative images of CD68^Lo^TMEM119^Hi^ and CD68^Hi^TMEM119^Lo^ microglia. (B-C) Quantification of homeostatic and phagocytic microglia in the ACC of the three groups. (n = 3-6 mice/sex/antibody, data shown are a percentage of 12 cells measured per animal across 3 images, one-way ANOVA, Sidak’s *post-hoc* test, main effect of antibody in legend). (D) Schematic of bulk RNASequencing of isolated microglia and whole forebrain. (E) PCA plot of isolated microglia transcriptomes from a-gp120 control and a-IL34 treated mice. (F and H) Volcano plot showing differentially expressed genes between a-gp120 control and a-IL34 treated microglia or whole forebrain (n=2-3 mice/sex/treatment, genes shown as significant passed a threshold of padj < 0.001 and LogFC > 1). (G and I) Quantification of microglia developmental index and LPS-induced “inflammatory” index from whole transcriptome data of isolated microglia or whole forebrain using methods described in Hanamsagar et al., 2017. (n=2-3 mice/sex/treatment, unpaired t-test). Related to Figure S5 and S7.

#### IL34 inhibition causes aberrant eating of VGlut2+ thalamocortical synapses

We next tested whether IL34-blocked microglia have elevated engulfment of VGlut2+ thalamocortical synapses. Consistent with the excitatory neuron-specific IL34 KO mice (Figure3), we found increased lysosomal content in anti-IL34 microglia and a two-fold increase in synaptic material contained within microglial lysosomes (Figures6A-C and S6B). Interestingly, there was a *decrease* in synaptic engulfment in anti-CSF1 mice, implying that blocking CSF1 has the opposite effect on microglia. We also found that anti-IL34 and anti-CSF1 have opposite effects on cell volume (anti-IL34 decreased volume while anti-CSF1 increased cell volume) (Figure S6C). There was a reduction in VGlut2+/PSD95 synapses in anti-IL34 treated mice, concordant with the synapse over-engulfment in these mice (Figures6D-E). Anti-CSF1 mice also have reduced synapse numbers, although this was presumably via a non-microglial mechanism since we observed decreased synaptic engulfment in these mice. The synapse decrease in anti-IL34 mice was driven by a reduction in VGlut2+ presynaptic puncta (Figure 6F). In sum, IL34 and CSF1 signaling have differential effects on microglia function at P15, and blocking IL34 increases thalamocortical synaptic engulfment at an inappropriate developmental window, leading to synapse loss We next tested whether these effects were generalizable to other IL34-dependent brain regions by performing similar analyses in hippocampal CA1 in our blocking antibody-treated mice (Figure 6G). Consistent with our findings in cortex, we saw no effect of anti-IL34 on microglia number in the hippocampus, but we did see a decrease in TMEM119 protein (Figures S6D-E). IMARIS reconstructions of hippocampal microglia revealed a significant increase in VGlut2 synaptic engulfment in anti-IL34 microglia, similar to cortical microglia (Figure 6H-I), despite no change in microglial volume or lysosomal content (Figures S6F-G). These data suggest that IL34 acts as a broad “don’t eat me” signal in both the ACC and hippocampus..

**Figure 6.**
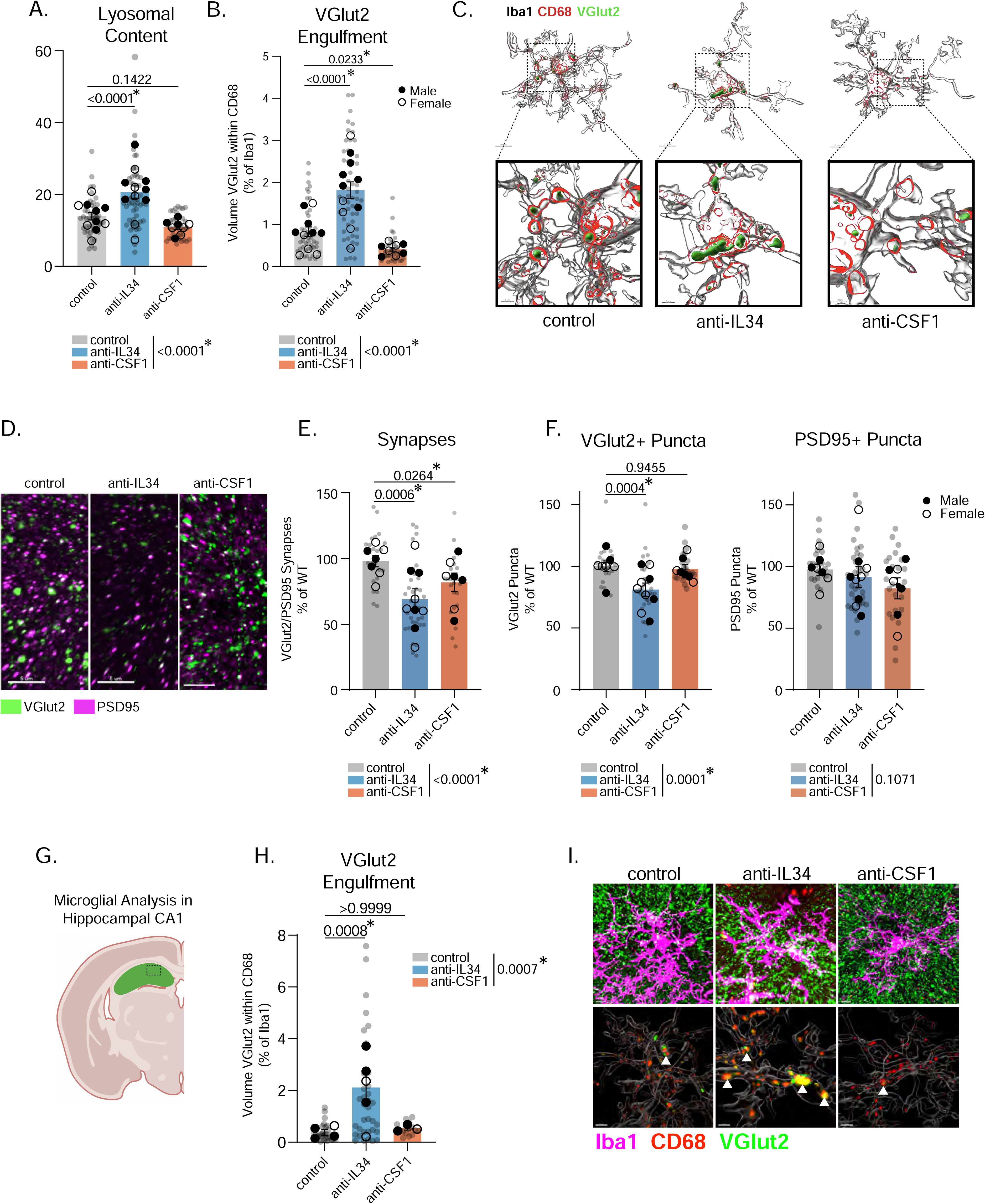
IL34 Inhibition Causes Aberrant Eating of VGlut2+ Synapses. (A-C) Quantification and representative images of lysosomal content and VGlut2 synaptic engulfment in control, anti-IL34, and anti-CSF1 treated mice. (n = 3-5 mice/sex/antibody, 4-6 cells analyzed per mouse, individual microglia represented by gray circles, animal averages represented by black dots, nested one-way ANOVA, Sidak’s *post-hoc* test, main effect of antibody in legend). (D-F) Representative images of VGlut2 and PSD95 and quantification of synapses in control, anti-IL34, or anti-CSF1 brains. (n = 3-5 litters/sex/antibody, 3 sections imaged per animal, 1-2 animals per litter, images represented by gray circles, litter averages represented by black dots, nested one-way ANOVA, Sidak’s *post-hoc* test, main effect of antibody in legend). (G) Schematic showing where hippocampal analysis was performed. (H-I) Quantification and IMARIS representative images of engulfed VGlut2 synaptic material in control, anti-IL34, and anti-CSF1 treated mice. (n = 1-3 mice/sex/antibody, 4-6 cells analyzed per mouse, individual microglia represented by gray circles, animal averages represented by black dots, nested one-way ANOVA, Sidak’s *post-hoc* test, main effect of antibody in legend). Related to Figure S6 and S7.

#### Acute IL34 inhibition does not cause mass microglia death and repopulation

To validate that the phagocytic, immature microglia in anti-IL34 mice were not newly born microglia repopulating after microglial cell death, we generated P2yr12^CreER/+^;tdtomato^fl/+^ mice to tag microglia before surgery. We injected 4-hydroxytamoxifen (4OHT) at P14, followed by ICV control or anti-IL34 antibodies at P15 and collected the mice 48 hours later (Figure S7A). We stained for Iba1 and RFP to determine what percentage of microglia after anti-IL34 were present before surgery and found >90% of Iba1+ cells were also RFP+ in both control and anti-IL34 mice (Figures S7B-C), confirming that the phagocytic microglia after a-IL34 are the same cells from before the antibody injection.

We next sought to confirm that the phagocytic microglia were not just the remaining microglia eating dying cells. There is evidence that astrocytes, not microglia, clear microglial debris following low-dose CSF1r-inhibition^39^. We injected control or anti-IL34 at P15 and collected mice 12- or 24-hours post-surgery. Interestingly, we observed an increase in microglia number in control mice 12 hours following surgery when compared to anti-IL34 mice or even control mice at 24- or 48-hours post-surgery (Figure S7D, dotted line). We believe that this is microglia proliferation occurring acutely after surgery caused by either physical stress or anesthesia (isoflurane) which is known to cause inflammatory responses in aged microglia^40^. We did not see this increase in microglia in anti-IL34 mice and only observed a slight reduction in microglia from baseline at 24 hours post-surgery. There was an increase in phagocytic cups (which are rare in P17 mouse cortex), in anti-IL34 mice at 12 hours post-surgery (Figures S7E-F). This corresponded to an increase in phagocytic, CD68^Hi^/TMEM^Lo^ microglia (Figures S7H and S7J), but no change in homeostatic, CD68^Lo^/TMEM^Hi^ microglia (Figures S7G).We believe this is due to a floor effect, as few control-treated microglia met the criteria for “homeostatic” this shortly after surgery. Finally, we observed a significant interaction between post-surgery time and antibody treatment in glial fibrillary acidic protein (GFAP) fluorescence demonstrating increased GFAP in anti-IL34 mice 24 hours post-surgery, which is temporally aligns with the slight decrease in microglia and suggests to us that astrocytes are eating any dying microglia (Figure S7I) in support of previous literature^39^, although we did not rigorously confirm this. In sum, our ICV injection causes microglial proliferation in control mice 12 hours post-surgery, an effect that is blunted by anti-IL34. Anti-IL34 increases phagocytic microglia 12 hours post-surgery, and any microglial death 24 hours post-surgery coincides with reactive astrocytes. We believe that these data confirm that the anti-IL34 microglia phenotypes are not a consequence of microglial engulfment of dying microglia immediately after surgery.

### IL34 overexpression at P1 increases TMEM119 and decreases synaptic engulfment at P8

Finally, we tested the sufficiency of IL34 signaling to control microglia maturation and synapse pruning. We took advantage of the fact that IL34 is expressed at low levels at P8 (Figures1 and 2) and virally overexpressed IL34 in neurons in the PFC of P1 mice (Figure 7A). This increased IL34 protein at P8 in the ACC (Figure 7A) and increased microglia number and TMEM119 protein (Figures7C-E). This is notable, as TMEM119 is not expressed in cortical microglia until the second postnatal week in mice^3^, suggesting that overexpressing IL34 before it is endogenously produced is, at least partially sufficient to accelerate the maturation of microglia. Next, we used IMARIS to measure lysosomal content and VGlut2 synaptic engulfment. Excitingly, IL34 overexpression reduced lysosomal content and thalamocortical synaptic engulfment (Figures7F-H and S7K), resulting in an increase in Vglut2+/PSD95 synapses (Figures7I-J). Finally, we observed a significant increase in microglial ramification in AAV-IL34 mice (Figure 7K).

**Figure 7.**
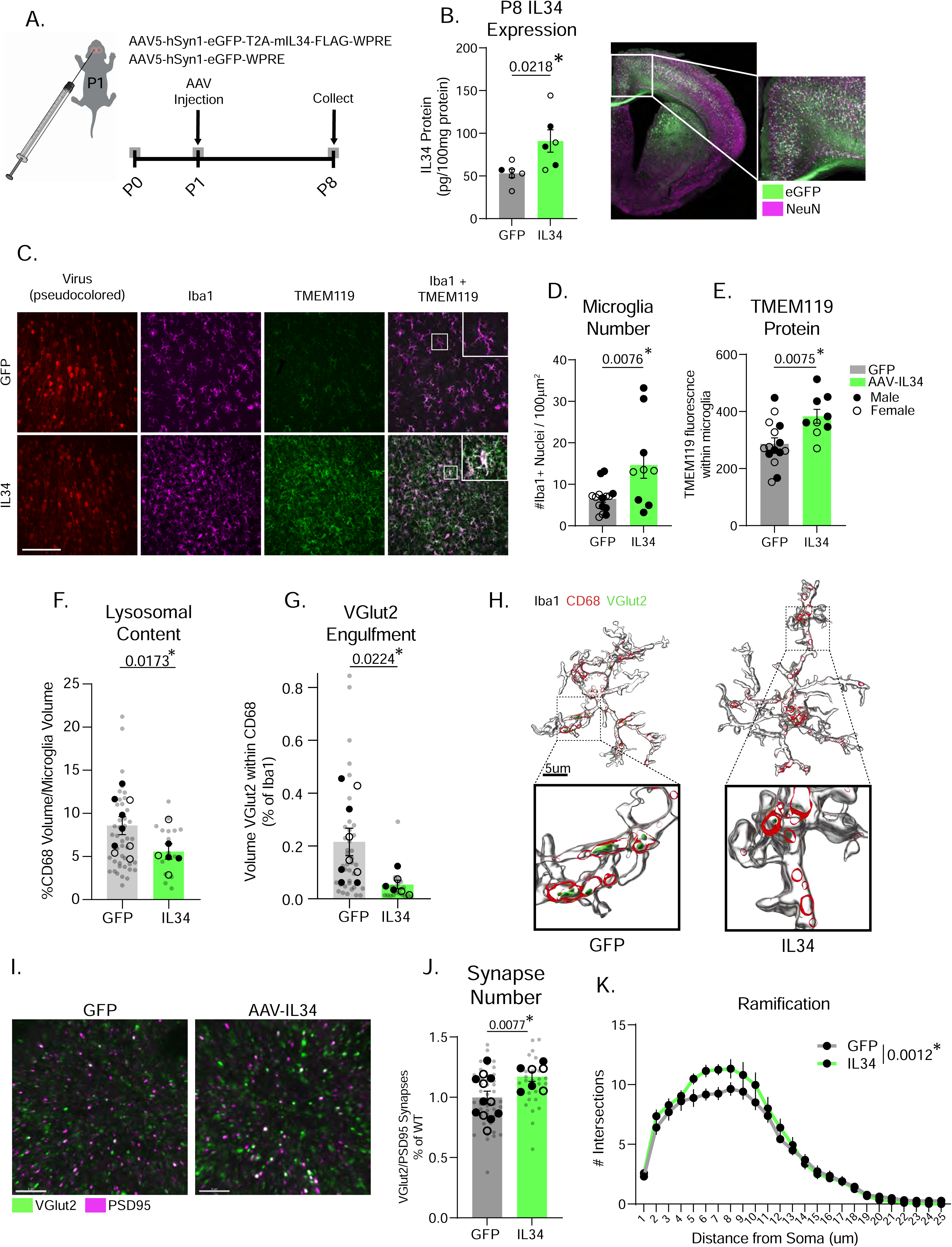
IL34 Viral Overexpression at P1 increases TMEM119 expression and reduces microglial engulfment of synapses. (A) Schematic of viral injections and tissue collection. (B) IL34 ELISA data measuring IL34 protein in control and AAV-IL34 mice. (n = 4-5 mice/group, unpaired t-test). (C-E) Representative images of Iba1 and TMEM119 and quantification of microglia numbers and TMEM119 in the ACC of mice injected with either control GFP or AAV-IL34. (n = 3-5 mice/sex/virus, unpaired t test). Scale = 100µm. (F) Quantification of microglia lysosomal content and VGlut2 synaptic engulfment and IMARIS representative images in control GFP and AAV-IL34 mice. (n = 3-5 mice/sex/virus, 4-6 cells analyzed per mouse, individual microglia represented by gray circles, animal averages represented by black dots, nested t-test). (I) Representative images and quantification of Vglut2+/PSD95+ overlap in control GFP or AAV-IL34 brains. (n = 3-4 litters/sex/antibody, 3 sections imaged per animal, 1-2 animals per litter, images represented by gray circles, litter averages represented by black dots, nested t-test). (K) Quantification of microglial ramification in control GFP and AAV-IL34 mice. (n = 3-5 mice/sex/virus, 4-6 cells analyzed per mouse, two-way ANOVA distance from soma x virus, p values shown are main effect of virus).

## Discussion

It is increasingly recognized that microglia exhibit a vast array of functional states throughout the developing brain^18,41–43^. We provide evidence that IL34, a neuron-derived cytokine that signals through the CSF1r on microglia, functionally matures cortical microglia in a discrete window of neurodevelopment. IL34 is upregulated in the second week of postnatal life and is expressed at a higher level in active, glutamatergic neurons in the ACC at P10. This developmental increase in IL34 expression in the ACC corresponds increased microglia numbers, ramification, TMEM119 expression, and a decrease in phagocytic capacity. Global genetic KO and neuron-specific KO of IL34 prevents these microglial developmental processes from progressing and causes overeating of excitatory synapses in the ACC. Acute blocking of IL34 at P15 increases immature, phagocytic microglia, decreases ramification and TMEM119 expression, and re-activates microglia to eat thalamocortical synapses during an inappropriate developmental window (P15). In turn, virally overexpressing IL34 in neurons early in development (P8) is sufficient to accelerate microglial functional maturity and inhibit synaptic pruning. These results establish IL34 as a “brake” for microglial phagocytosis in the context of synaptic pruning, as well as unveiling its greater role in fine-tuning microglial identity and functional state in development.

We found that IL34 expression is higher in Fos+, glutamatergic neurons. Despite these exciting initial findings, follow up work is needed to define its release dynamics from neurons. We were limited with our RNAFISH experiments to assessing mRNA only within the nucleus, but there is transcriptomic data suggesting that IL34 mRNA is present at the synapse and may be locally translated there^44^. This is relevant since our finding that VGlut2-specific KO of IL34, which results in the loss of IL34 in excitatory thalamic projection neurons, was sufficient to reduce microglia numbers and increase engulfment of synapses in the cortex, suggesting the potential for thalamic neurons to influence cortical microglia via synaptic IL34 release. Additionally, we found that blocking IL34 specifically increased microglia engulfment of presynaptic (VGlut2+) terminals in the ACC. This does not, however, inform what role it may play in engulfing post-synaptic material, which microglia have been shown to do^45^, or other neuronal components such as spines or cell bodies in other brain regions. As a secreted protein, it is unlikely that IL34 is tagging individual synapses for engulfment like complement^22^. Thus, based on our data, we believe that IL34 has a broad effect on local microglia once the circuit has stabilized. The discovery that *IL34* gene expression is lower in GABAergic neurons compared to glutamatergic neurons at postnatal day 10 was not surprising to us. While the timing of putative inhibitory synaptic pruning in the ACC is not well described, GABAergic pruning generally occurs later in development than excitatory synaptic pruning^46^, in part due to the delayed maturation of GABA circuits as the GABA switch happens in the second postnatal week^47^. It stands to reason that GABAergic neurons may upregulate IL34 expression directly following their circuit maturation, and this should be addressed in future work.

While both IL34 and CSF1 bind to the same receptor on microglia, our findings demonstrate that they have distinct effects on microglial functional differentiation, with separate roles in development. Prior to our work, *Kana et al.*, demonstrated that microglia in culture treated with IL34 vs. CSF1 have different transcriptional responses, but no data existed that tested the role of these two molecules on microglia function in an intact, developing brain^14,48^. IL34 binding to CSF1r in macrophages leads to increased phosphorylation of downstream targets in the MAPK pathway, while CSF1-binding preferentially phosphorylates members of the STAT and ribosomal S6K pathways^49^, which may provide one explanation for their differential effects on cell function. Additionally, IL34 polarizes macrophages to a less “activated” state, by increasing IL-10 and decreasing phagocytosis *in vitro*^50^, and microglia in gray matter are sensitive to IL34 blocking or ablation, while microglia in white matter, cerebellum, and brainstem are sensitive to CSF1 blocking or ablation^12,13^. This is consistent with the finding that white matter microglia exhibit different functional and transcriptomic properties compared to gray matter microglia^51,52^.Our experiments demonstrate that IL34 signaling is critical not only for gray matter microglial survival and proliferation at these early stages of life, but also for dictating the proper functional development of these cells.

Interestingly, microglia number and TMEM119 in the cortex of IL34 KO mice were indistinguishable from cerebellar microglia, suggesting CSF1 maintains a subset of microglia in the cortex when the primary signal (IL34) is lost^5,11^. The question remains, however, whether two distinct populations of microglia are maintained solely by IL34 or CSF1 in the intact brain. This is unlikely given that CSF1 is still expressed at low levels in the forebrain of wild type mice^13^. Thus, we hypothesize that IL34/CSF1 functional regulation exists along a spectrum – where the balance of total IL34 vs. CSF1 present in the system can tip cells back and forth between various states. This suggests that IL34/CSF1 signaling can differentially modulate microglial function across various developmental and disease states and warrants future studies that interrogate the potential role for IL34 signaling in diseases where microglia play a causal role.

Intriguingly, a stop-gain mutation in the IL34 gene may confer risk for Alzheimer’s disease (AD) in humans^53,54^. Further, IL34 expression levels are decreased (while CSF1 levels are increased) in Alzheimer’s disease and in animal models^55,56^, and IL34 infusion is sufficient to rescue associative learning deficits in APP/PS1 mice^57^. In addition to the potential implications for AD, an abnormal balance in IL34 and CSF1 signaling has been demonstrated in several neurodegenerative diseases, including Huntington’s, multiple sclerosis, amyotrophic lateral sclerosis, and chronic neuropathic pain^58–62^. The basic biological mechanisms uncovered from our studies provide a foundation for future experiments to interrogate the functional role of IL34 signaling in these conditions and position it as a potential therapeutic target in a range of degenerative conditions.

We observed an unexpected decrease in synapses in the anti-CSF1 mice, despite a decrease in microglial engulfment of pre-synaptic structures compared to control-treated mice (Figure 6). This suggests that CSF1 may be important in other aspects of synaptic development. It is possible that our anti-CSF1 manipulation is directly impacting neurons, since there is evidence that neurons can express the CSF1 receptor and that signaling through this receptor is protective^35^, however we did not rigorously test this hypothesis. There are other reports suggesting CSF1 is produced by neurons, but only in the context of stress^63^, injury^58^, or ethanol withdrawal^64^, and that this upregulation has negative consequences for neurons and synaptic transmission. These findings are in line with our observations that blocking CSF1 decreased microglial engulfment of synapses and suggest that CSF1 may be primarily involved in maintaining immature phagocytic microglia. More work is needed to tease out the precise effects of CSF1 and IL34 signaling in different cell types in different contexts. Our data also demonstrate a need for greater attention to and interpretation of results utilizing CSF1r inhibitors. It sets a precedent for modeling different microglial functional states *in vitro* using different growth factors (CSF1 vs. IL34) and demands additional scrutiny of these models for replicating an intact brain environment (with homeostatic levels of these cytokines across contexts)^65^.

To summarize, our results unveil a mechanism by which neurons communicate with microglia to control their function in postnatal development. In addition to promoting survival, IL34 signaling to microglia acts as an important cue that matures microglia and prevents over-eating of thalamocortical synapses. These data shed light on the functional relevance of IL34, making it a unique and interesting target for future studies of microglia and neurons.

### Limitations of the study

While our blocking antibody approach complemented our genetic KO data and prevented significant microglia cell loss, there were some limitations in that the manipulation lacked cell-type specificity - it blocked binding of all cellular sources of IL34 and was not specific to microglia. Additionally, perhaps the largest limitation of our study was that we have yet to identify the mechanism by which IL34 signaling programs microglia function. While we expect that the CSF1 receptor is involved, we were unable to empirically test this because of the receptor’s necessary role in microglial survival. Further, questions remain regarding how and why IL34-deficient microglia overeat excitatory synapses. We hypothesize that it is by programing microglia to change their “sensitivity” to other, more specific signals including: C3/C1q, Sirpa/CD47, CX3CR1/CX3CL1, Adam10, or Trem2/externalized phosphatidylserine^22,27,29,31,32,66^ however we did not experimentally confirm this. Finally, we did not overexpress CSF1, so we are unable to confirm that CSF1 overexpression in a cell type where CSF1 is not normally expressed (neurons) would have a different effect.

## Resource Availability

### Lead Contact

Requests for further information and resources should be directed to and will be fulfilled by the lead contact, Dr. Staci Bilbo.

## Materials Availability

This study did not generate new unique reagents.

## Data and code availability

All original code for analysis is available at https://github.com/bendevlin18 and bulk RNASequencing data generated is deposited in GEO: accession GSE290856. Any additional information required to reanalyze the data reported in this paper is available from the lead contact upon request.

## Acknowledgements

This work was supported by funding from the NIH (1F31NS130757-01), the Cure Alzheimer’s fund to B.A.D and S.D.B, and NIDA-DA047233, NINDS-R01NS106721, NIA-R01AG072489, ERC-951515 awarded to A.S. We would like to thank D. Saban for the IL34^LacZ/LacZ^ mice, R. Weimer from Genentech for the IL34 and CSF1 function-blocking antibodies, M. Alter for his continuous and invaluable discussion of the work, S. Monroe for their help with the graphical abstract, and the Duke Light Microscopy Core facility for microscope and image analysis support.

## Author Contributions

S.D.B and A.M.C conceived the study and, together with B.A.D, designed the experiments. B.A.D, D.M.N, D.R., G.G., M.J.C, S.O., M.D., A.S., S.A., and A.F. performed experiments; D.R. and A.S. provided critical reagents, B.A.D analyzed all data and wrote the manuscript together with S.D.B, who oversaw the project.

## Inclusion and Diversity

We support inclusive, diverse, and equitable conduct of research.

## Main Tables and Legends

No tables are included in this manuscript.

## STAR Methods

### Animals

All procedures relating to animal care and treatment conformed to Duke Institutional Animal Care and Use Committee and National Institutes of Health guidelines (protocol no. A107-19-05 and renewal no. A062-22-03). Animals were group housed in a standard 12:12-hour light–dark cycle. The C57BL/6J mouse line was obtained from Jackson Laboratory, stock no. 000664. The IL34^LacZ/LacZ^ mice were a gift from Dr. Daniel Saban at Duke University and were originally generated by Dr. Marco Colonna^10^.

VGlut1-Cre male mice (B6.Cg-Slc17a7^tm1.1(cre)Hze/J^) were ordered from Jackson Laboratory, stock no. 037512 and bred in house with C57BL/6J females to produce VGlut1-Cre offspring for cre-dependent chemogenetics experiments. VGlut2-Cre mice (B6J.129S6(FVB)-Slc17a6^tm2(cre)Lowl^/MwarJ) were ordered from Jackson Laboratory, stock no. 028863, and bred in house with IL34^fl/fl^ mice (generous gift from Dr. Marco Colonna to Dr. Anne Schaefer^10,13^) to produce VGlut2-Cre;IL34^fl/fl^ offspring for excitatory neuron-specific IL34 KO experiments. Ai14 tdtomato^fl/fl^ mice were ordered as breeder females from Jackson Laboratory, stock no. 007914 and B6(129S6)-P2ry12^em1(icre/ERT2)Tda^/J (P2yr12Cre^ER/+^) mice were recently restored from cryopreservation by the INIA (Integrative Neuroscience Initiative on Alcoholism) consortium and were gifted to S.D.B. These mice are now available from Jackson Laboratory, stock no. 034727.

#### qPCR

For the qPCR time course, wild-type mice were sacrificed and saline perfused at the six described ages (P7, P14, P21, P30, P38, P55) and their brains flash-frozen in 2-methylbutane in dry ice and stored at -80C. Tissue punches were taken from the anterior cingulate cortex, nucleus accumbens, amygdala, and cerebellum and stored in 500µL TRIzol (ThermoFisher, 15596026). RNA was extracted from the tissue using TRIzol based chloroform extraction, followed by isopropanol precipitation. Resulting RNA concentration was measured using a NanoDrop spectrophotometer (ND- 1000) and either 200 ng or 1000 ng (depending on starting concentration) of RNA were reverse transcribed into cDNA using a Qiagen QuantiTect Reverse Transcription kit (205311). qPCR was performed on an Eppendorf Realplex ep Mastercycler using Sybr/Rox amplification with a QuantiFAST PCR kit (204056). Primer sequences are listed in Supplementary Table 1. All samples were loaded in triplicate and all plates were run on the same day to ensure comparability across plates. Fold change was calculated using the 2^−ΔΔCT^ method with 18S as an endogenous control for sample normalization.

#### IL34 ELISA

For all ELISA tissue collection, mice were anaesthetized with CO_2_ and transcardially perfused with ice-cold saline prior to brain collection. For tissue punches, brains were flash-frozen in dry ice and embedded in O.C.T. (Sakura Finetek) and stored at -80C. Tissue was sectioned on the Leica CM50 cryostat until brain region of interest was visible, then brain region was punched using a 1mm RapidCore tissue punch (Ted Pella) and frozen at -80C. Prior to the tissue-punch ELISA, tissue was homogenized in 300µL 1X PBS using the Dremel Tissue-Tearor (BioSpec, 985370-04). Then, 300µL Cell Lysis Buffer 2 was added to the samples (R&D Biosystems, 895347) and they were placed on an orbital shaker at 200rpm for 30 minutes before spinning down in a 4`°`C centrifuge at 13,200 rpm for 20 minutes. Samples were stored at -20C overnight or -80C long term. All samples were run according to the manufacturer’s instructions for the kit (R&D Biosystems, M3400). All protein was normalized to total protein in the sample using a standard bradford assay kit (BioRad, DC™ Protein Assay Kit, 5000111), which was assessed the same day as the ELISA. The equation used to calculate pg/100mg protein is: (IL34 concentration (pg/mL) *100) / (total protein (mg/mL) *1000).

#### Microglia Staining

For all immunohistochemistry experiments, mice were perfused with ice-cold saline followed by 4% paraformaldehyde (PFA). Tissue was then post-fixed in 4% PFA for 24 hours prior to cryoprotection in 30% sucrose with 0.1% sodium azide. Brains were flash-frozen in 2-methylbutane in dry ice and stored at -80C until sectioned. Sections were collected at 40µM and kept free-floating at -20C in cryoprotectant until stained. Free-floating sections were washed 3X in 1X PBS prior to blocking in a 10% normal goat serum (NGS) solution with 0.3% Triton-X in 1X PBS for 1 hour. Sections were incubated in primary antibodies overnight at 4`°`C. The primary antibodies used were Chicken anti-Iba1 (Synaptic Systems, 234 009, 1:1000), Guinea Pig anti-TMEM119 (Synaptic Systems, 400 004, 1:1000), Rat anti-CD68 (Biolegend, 137002, 1:1000). The following day, sections were washed 3X in 1X PBS and incubated in secondary antibodies for 2 hours at room temperature prior to mounting on gelatin-subbed slides and cover-slipping with Vectashield plus DAPI (Vector Labs, H-2000-10).

#### Microglia Imaging and Analysis

Images of microglial Iba1, TMEM119, and CD68 staining were acquired on a Zeiss AxioImager M2 at 20X magnification. 19 Z-Stacks at a 1µM step size were acquired and maximum intensity projections were used for all analyses. For *Microglia Cell Counting*, single channel, maximum intensity, Iba1 images were counted using the supervised machine learning tool Ilastik^67^ with the Pixel + Object Segmentation pipeline. All automatic counts were validated by blinded hand counts. Object segmentation outputs were then combined and analyzed using custom python scripts. For *Microglial TMEM119 expression,* first, an Iba1 mask was generated using Ilastik’s pixel segmentation pipeline on single channel Iba1 images. This mask was used to control for any mean gray value differences that may be a result of decreased cell number or process complexity. Following mask generation, TMEM119 mean gray value within the Iba1 mask was calculated using a custom Fiji script. All details and code for these analyses can be found at github.com/bendevlin18/mgla_img_analysis_pipeline. Finally, for *Microglia 2D Sholl Analysis*, using the Iba1 stain, individual microglia were selected for analysis from each image by an individual blind to sex, genotype, and age. Those individual microglia were put through the Ilastik pixel segmentation pipeline to generate binary segmentations of the images. These binary segmentations were processed using a custom python script based on the sholl analysis fiji plugin that skeletonizes the image and plots concentric rings from the soma of each cell, measuring the intersections of those rings with cell processes. All code for 2D sholl analysis is also available on GitHub. https://github.com/bendevlin18/sholl-analysis-python.

#### Microglia Phagocytic Capacity Imaging and Analysis

For microglia phagocytic capacity, microglia (Iba1) and CD68 staining was imaged using a Leica SP8 upright confocal. Individual microglia were imaged at a 63X magnification with a Z-stack step size of 0.33 µM. These images were then imported into IMARIS (Oxford Instruments, v9.0) and surfaces rendered of the Iba1 and CD68 channels to generate 3D reconstructions of the microglia and their lysosomes. The phagocytic capacity calculation was as follows: (total CD68 volume within microglia / total microglial volume (Iba1)) *100.

#### Microglia Heterogeneity Quantification

To quantify microglial heterogeneity, representative microglia were selected by a blind experimenter from every image using the Iba1 channel. Each cell was selected using the freehand selection tool in Fiji (ImageJ) and TMEM119 and CD68 mean gray value were measured within each selection. 4 cells were measured per 20X images, and 3 images were analyzed per animal. TMEM119 and CD68 measurements were then normalized to Iba1 expression and histograms representing the distribution of expression values were plotted to determine value cutoffs for high, low, and mid expressing cells of each marker (supp. Fig. 7b-c). Individual cells were assigned high, low, or mid based on these cutoffs, and percentages of cells (out of 12 total) falling into each category were calculated per animal. The entire python script used for this analysis is available on GitHub. https://github.com/bendevlin18/microglia_tmem_cd68_heterogeneity.

#### RNA Sequencing

Mice were saline-perfused, and their brains dissected and split at the midline using a razor blade. Whole forebrain was collected, and microglia were isolated from one half using a CD11b antibody-based procedure, according to published methods^68^. RNA was extracted from whole forebrain and CD11b+ isolated microglia using the TRIzol based chloroform extraction and samples were transferred to MedGenome (Foster City, California) for library preparation and sequencing. Raw fastQ files were aligned to Mus Musculus GRCm38 mm9 using STAR (v2.7.5c) and featurecounts (v1.6.3) on the Duke Compute Cluster with custom bash scripts. Genes were filtered and only included in analysis if they were present at 10 counts in at least 4 samples. Differentially expressed genes were calculated using DESeq2^69^ in R 4.4.0. All analysis scripts are available on GitHub. https://github.com/bendevlin18/IL34blocking_seq_analysis_2024.git.

#### Neuron, Astrocyte, Oligodendrocyte Stain and Quantification

Free-floating sections were washed 3X in 1X PBS prior to blocking in a 10% normal goat serum (NGS) solution with 0.3% Triton-X in 1X PBS for 1 hour. Sections were incubated in primary antibodies overnight at 4`°`C. The primary antibodies used were Rabbit anti-Sox9 (Millipore, AB5535, 1:1000), Guinea Pig anti-Neun (Synaptic Systems, 266 004, 1:1000), Mouse anti-Olig2 (Millipore, MABN50, 1:250). The following day, sections were washed 3X in 1X PBS and incubated in secondary antibodies (1:200) for 2 hours at room temperature prior to mounting on gelatin-subbed slides and cover-slipping with Vectashield plus DAPI (Vector Labs, H-2000-10). To quantify overall numbers of neurons, astrocytes, and oligodendrocytes in the cortex and braitem, images were acquired on a Zeiss AxioImager M2 at 20X magnification. 19 Z-Stacks at a 1µM step size were taken and maximum intensity projections were used for all analyses. Single channel, maximum intensity, images of each of the three stains were counted using Ilastik with the Pixel + Object Segmentation pipeline. All automatic counts were validated by blinded hand counts. Object segmentation outputs were then combined and analyzed using custom python scripts.

#### Synaptic Staining

Samples were collected and processed as described above. Free-floating sections were rinsed in 1X TBST (0.2% Triton X-100) 3 times for 10 minutes prior to block. Sections were then incubated in 5% NGS in 1X TBST for 1 hour at room temperature. Sections were then incubated in primary antibody diluted in 5% NGS in 1X TBST overnight at 4`°`C. The primary antibodies used were Guinea Pig anti- VGlut2 (Synaptic Systems, 135 404, 1:2000), Rabbit anti-PSD95 (ThermoFisher, 51-6900, 1:350) for overall synapse counts, or Guinea Pig anti-VGlut2 (Synaptic Systems, 135 404, 1:2000), Rat anti- CD68 (Biolegend, 137002, 1:1000), and Chicken anti-Iba1 (Synaptic Systems, 234 009, 1:500). Following primary incubation, sections were washed 3X for 10 minutes in TBS and then incubated in secondary antibody diluted in 5% NGS in 1X TBST for 2 hours at room temperature. Sections were mounted on gelatin-subbed slides and cover-slipped with Fluoromount G (ThermoFisher, 00-4959- 52).

#### Synaptic Imaging and Analysis

Synapse images were acquired on an Olympus FV3000 inverted confocal. Images were taken at 60X magnification with a 1.6X optical zoom. A 4x4 tile scan with 4, 0.33 µM step Z-stacks were obtained. Synapse images were analyzed using SynBot^70^. Synapses were identified by the colocalization of pre- (Vglut2) and post- (PSD95) synaptic puncta.

#### Synaptic Engulfment Imaging and Analysis

Synaptic Engulfment images were acquired on an Olympus FV3000 inverted confocal. Images were taken at 60X magnification with a 2.3X optical zoom. A 2x2 tile scan with 100-110, 0.33 µM step Z- stacks were obtained. Whole, stitched image stacks were imported into IMARIS (Oxford Instruments, v9.0) and microglial surfaces were rendered using the Iba1 channel. Individual microglia were selected and reconstructed (up to 3/image) and used to mask the CD68 channel. A surface was created from each Iba1-masked CD68 channel and the CD68 surface was used to mask the VGlut2 channel. Surfaces were created of the CD68-masked VGlut2 signal and volumes of all surfaces were exported for analysis. The phagocytic capacity was calculated as described above. The volume of synaptic material engulfed was normalized to Iba1 cell volume.

### Behavior

#### Ultrasonic Vocalizations

We performed ultrasonic vocalization testing as previously described^25^. Briefly, pups were removed from their cage and individually placed in a cotton-lined cup within a sound-attenuating chamber under an Avisoft Condenser ultrasound microphone (Avisoft Bioacoustics CM16/CMPA). suspended four inches above the contained pup. The dam and any remaining littermates were removed from the testing room in the home cage. After 3 min of recording, pups were weighed and sexed before being returned to their home cage. After the dam attended to each returned pup, the home cage was returned to the colony room. Analysis of USV .wav files was done using the MATLAB (MatlabR_2022a) machine-learning program MUPET to quantify the number of calls, the energy of the calls, and all syllable information. Forty-unit syllable repertoires were generated for each dataset and manually inspected before USV similarity was calculated and heatmaps were generated.

#### Open Field

Adult mice (P50-55) were placed in a 50 cm × 50 cm square enclosure with 38-cm-high walls and allowed to explore freely for 10 minutes. Behaviors (distance moved and velocity) were recorded and analyzed using EthoVision video tracking software (Noldus). Center avoidance was assessed by comparing time spent at the periphery of the chamber to time spent in the center (middle third of chamber).

#### Social Preference

Adult mice (P60-65) were assessed for social preference using a three-chambered preference test. On test day, subject and stimulus animals were habituated to the testing room for a minimum of for 1 h before testing. Each mouse was then tested for the preference to investigate a novel object (rubber duck) versus a novel social stimulus (age and sex-matched stimulus animal). Mice were placed in the middle chamber with the rubber duck confined in a clear plexiglass cylindrical cage on one side of the test and the novel conspecific confined in an identical cage on the other side for 10 minutes. The test was recorded with a Logitech webcam, and all behavior was manually quantified using Solomon Coder by an observer blind to sex and treatment. The social preference score was calculated as: (time spent investigating social stimulus / total time spent investigating either stimulus) * 100.

#### Elevated Plus Maze

Male and Female adult mice (P70-75) were tested individually in the elevated plus maze (each arm 10cm wide by 52 cm long) to measure anxiety-like behavior. On the day of testing, mice were habituated to the testing room for at least 1 hour prior to start. Males and females were tested on separate days. One by one, each mouse was placed in the center of the elevated plus maze and their movement throughout the maze was tracked for 5 minutes. Behaviors were recorded and analyzed using Ethovision (Noldus), and the endpoints measured are as follows: Distance Moved, Velocity, Time in Open Arms, Time in Closed Arms, Time in Center, # of Transitions from Open to Closed Arms. The “Percent in Open Arms” metric was calculated by (time in open arms / time in closed arms) *100. *<u>Barnes Maze</u>*.

Male adult mice (P80-90) were tested for memory/cognitive flexibility on Barnes maze over a 5-day period. In the first 3 days, each mouse was placed in the center of a large (45cm, radius) circular platform and given 4 minutes to find and enter the target hole (1 of 20 holes with an escape hatch). This was repeated a total of 3 times for each mouse on each of the first 3 days (Training Phase), and the amount of time taken to find the target hole drastically decreased across the 3 days, indicating that the mice had learned the location of the target hole. On the fourth day, the target hole was moved to another quadrant of the maze, and the mice were tested for 3 trials on day 4 and 5 to measure their reversal learning potential. Ethovision (Noldus) was used to track their performance in this task, and the endpoints analyzed were Time to Escape, Total Nose poke Errors, Time Spent in Original Target Quadrant (reversal only).

#### P1 Viral Injections

Postnatal day 1 mouse pups were placed in a 15mL conical tube and anesthetized in an ice bath for 10 minutes. Once pups were unresponsive via toe pinch, they were removed from the conical tube and placed on an ice pack for the duration of the injections. Mice were injected with a glass pulled needle pre-filled with virus connected to a microinjector (KD Scientific, 788100). 200nL was injected over a 30 second period into each hemisphere, and prefrontal cortex was targeted by injecting 2mm posterior to the inferior cerebral vein and 1mm lateral to the sagittal sinus. For the *DREADDs* experiments, we injected one of the following viruses: AAV8-hSyn-hM3D(Gq)-mCherry (Addgene, 50474-AAV8), AAV8- hSyn-hM4D(Gi)-mCherry (Addgene, 50475-AAV8), or AAV8-hSyn-EGFP (Addgene, 50465). For the *IL34 Overexpression* experiments, we injected one of the following custom-made viruses: AAV5-hSYN1-eGFP-WPRE (control) or AAV5-hSYN-eGFP-T2A-mIL34-FLAG-WPRE (IL34 overexpressing).

#### P15 Blocking Antibody Injections

Mice were anesthetized using 4% isoflurane pretreated with Ketophen anesthetic (5mg/kg, S.C.), and fitted into a Stoelting stereotaxic apparatus for the duration of the surgery. The fur on the head was shaved and, following betadine and ethanol preparation of the skin, a single incision was made. The surface of the skull was cleaned and Bregma was identified. All blocking antibodies were obtained from Genentech^12^, stored at 4`°`C and diluted to a working stock of 10mg/mL in sterile saline on the day of surgery. A 10uL 1701RN Hamilton syringe (Thomas Scientific, 1207K95) with attached pulled glass needle was loaded with 4µL of antibody and lowered into each ventricle (-.7AP, 1ML, 2.4DV to bregma). 2µL were diffused over 2 minutes into each ventricle.

#### Microglia P2RY12^CreER^ experiments

Mice were injected IP with a single dose of 75mg/kg 4OHT or saline dissolved in corn oil at P14 and underwent blocking antibody surgery the following day at P15. Surgeries were performed in the same manner as all other blocking antibody experiments and mice were collected 2 days after surgery at P17.

#### DREADD DCZ Injections

DCZ (Tocris, 7193) was dissolved in 1% DMSO in sterile saline at a working dilution of 0.05 mg/mL. For the neonatal (P8-10) DREADD DCZ injections, mice were weighed daily prior to injection and injected with a 1mg/kg dose of DCZ at the same time each day (11a.m.). The mice were collected 2 hours following DCZ administration on the third day of injections (P10). For the adult DREADD DCZ injections, mice were weighed and injected 15 minutes apart to allow for collection 90 minutes post DCZ administration.

### RNA In Situ Staining, Imaging, and Analysis

#### RNAScope

Brains for RNAScope were collected 2 hours following the final DCZ injection at P10. The mice were deeply anesthetized using Avertin (tribromoethanol) and transcardially perfused with ice-cold saline and their brains embedded in O.C.T. (Sakura Finetek) and flash frozen in dry ice. The brains were sectioned at 20uM thickness and thaw mounted onto Superfrost Plus slides (Fisher Scientific, 12-550- 15). Slides were stored at -80C until stained. We performed RNAScope (ACDBio) in accordance with the manufacturer’s protocol. Briefly, tissue was fixed for 15 minutes in 4% PFA followed by gradual dehydration for 5 minutes each in 50%, 70% and 100% EtOH solutions. The samples were then treated with protease for 30 minutes, briefly washed, and then incubated with primary probes for 2 hours at 37`°`C. The primary probes used were C1-Gad2, C2-VGlut1, and C3-IL34. Following hybridization, the tissue went through a series of 4 amplification steps per kit instructions. After the fourth amplification, the tissue was washed, and coverslipped with Vectashield plus DAPI (Vector Labs, H-2000-10). Slides were stained in batches of 4 and imaged within 72 hours of the staining on a Leica SP8 upright confocal. 19 step Z-Stack images at a thickness of 0.33µM of the ACC were taken with the 40X objective. 3 sections per animal were imaged and included in the analysis.

#### HCR RNAFish

Brains for HCR RNAFish (Molecular Instruments) were collected and sectioned and pretreated (fixation and dehydration) in the same manner as RNAScope. Following fixation and dehydration, the tissue was placed in primary probe solution overnight at 37`°`C. The primary probes used were B1- Gad2, B2-Fos, and B3-IL34. The next day, tissue was washed at 37`°`C in increasing concentrations of 5X SSCT buffer and then incubated with hairpins directed against the primary probes overnight at 37`°`C. On the final day, the tissue was washed in 100% 5X SSCT and cover slipped using Fluoromount G (ThermoFisher, 00-4959-52). Imaging of these samples was performed using an Olympus Fluoview 3000 inverted confocal with the 30X objective on the same settings as the RNAScope imaging.

#### Analysis

Images from RNAScope and RNAFish pipelines were analyzed in IMARIS (Oxford Instruments, v9.0) using the “Imaris for Cell Biologists” plugin to generate cells from the DAPI stain and to count each RNAScope puncta as a separate “vesicle” type within that cell. The volume of the DAPI stain was included in the output as well as the number of each vesicle type. Every cell in the image was included in the analysis and cells were determined to be “glutamatergic” or “GABAergic” based on a threshold adjusted number of Vglut1 puncta vs. Gad2 puncta. Additionally, cells were determined to be Fos+ or Fos- based on an average of 6 puncta adjusted for cell size. IL34 puncta number was normalized to total DAPI volume and averaged for all cells of a given type within a given animal. The code used to process the data is publicly available at https://github.com/bendevlin18/RNA_in_situ_single_cell_quant.git.

#### Statistics

Statistical tests were performed using GraphPad Prism 9. Raw data as well as a description of the tests and results (including multiple comparison corrections and post-hoc analyses) are provided. In all experiments, both male and female mice were included at sufficient power (n ≥ 3 biological replicates) and statistics were first run including sex as a variable (e.g. three-way ANOVA with sex x age x genotype). In all cases where there was not a main effect of sex, male and female mice were then combined for subsequent statistical testing. Unless otherwise noted, data are mean ± S.E.M. and all p-values that fall below p<0.05 are denoted with an asterisk, while p-values between 0.05 and 0.10 denoted with a #.

---

**Supplemental Figure 1.**
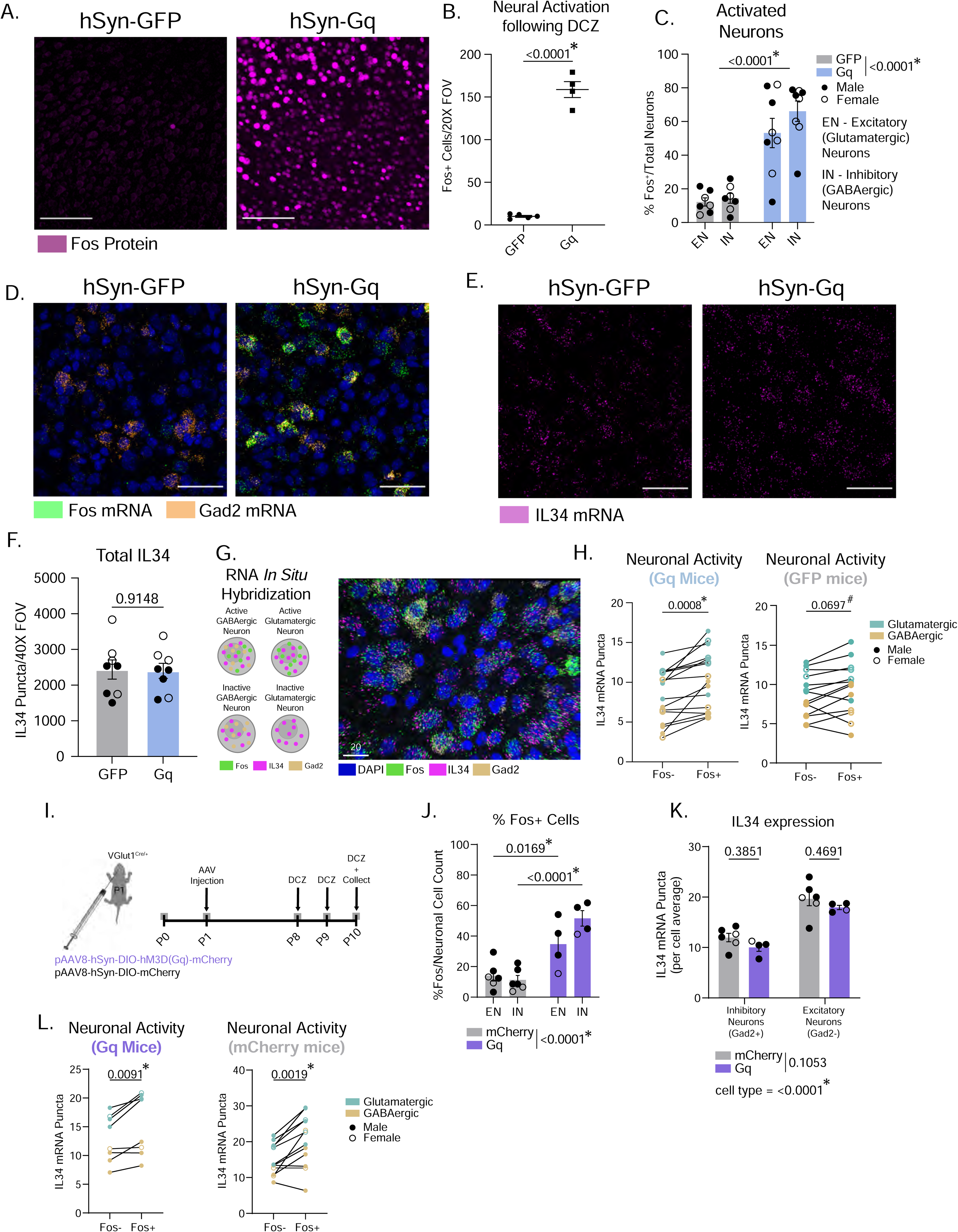
IL34 expression is increased in Fos+ neurons. Related to Figure 1. (A-B) Representative images and quantification of Fos protein in GFP-control mice and Gq-DREADD mice. (n = 4-5 mice/virus, unpaired t-test). Scale = 100uM (C-D) Quantification and representative images of Fos mRNA in excitatory and inhibitory neurons in GFP and Gq mice. (n=3-4 mice/sex/virus, one-way ANOVA, main effect of virus in legend). Scale = 50uM (E-F) Total IL34 puncta per 40X field of view in GFP control and Gq mice. (n=3-4 mice/sex/virus, unpaired t-test). Scale = 50uM (D) Schematic outlining the different cell types identified by expression of IL34, Gad2, and Fos using RNA-FISH and representative image of RNA-FISH stain showing IL34 levels in Fos+ (active) and Fos- (inactive) neurons. (E) Quantification of IL34 mRNA puncta in Fos- and Fos+ cells in mice that received the Gq or GFP virus. (n = 4 mice/sex, data shown are a per animal average of the expression level of all neurons of that type (excitatory vs. inhibitory, and active vs. inactive, total of 5,722 cells analyzed), paired t-test). (F) Schematic of excitatory neuron-specific chemogenetic activation experiments. (G) Quantification of % Fos+ cells in excitatory and inhibitory neurons in mCherry and Gq mice. (n=1-4 mice/sex/virus, data shown are an average of 3 images taken from 3 sections of each mouse, two-way ANOVA, Sidak’s post-hoc test, main effect of virus shown in legend). (H) Quantification of average IL34 expression in excitatory and inhibitory neurons in mCherry and Gq mice. (n=1-4 mice/sex/virus, data shown are an average of 3 images taken from 3 sections of each mouse, two-way ANOVA, Sidak’s post-hoc test, main effect of virus in legend). (I) Quantification of IL34 mRNA puncta in Fos- and Fos+ cells in mice that received the Gq or control virus. (n=1-4 mice/sex/virus, data shown are a per animal average of the expression level of all neurons of that type (excitatory vs. inhibitory, and active vs. inactive, total of 2,566 cells analyzed), paired t-test).

**Supplemental Figure 2.**
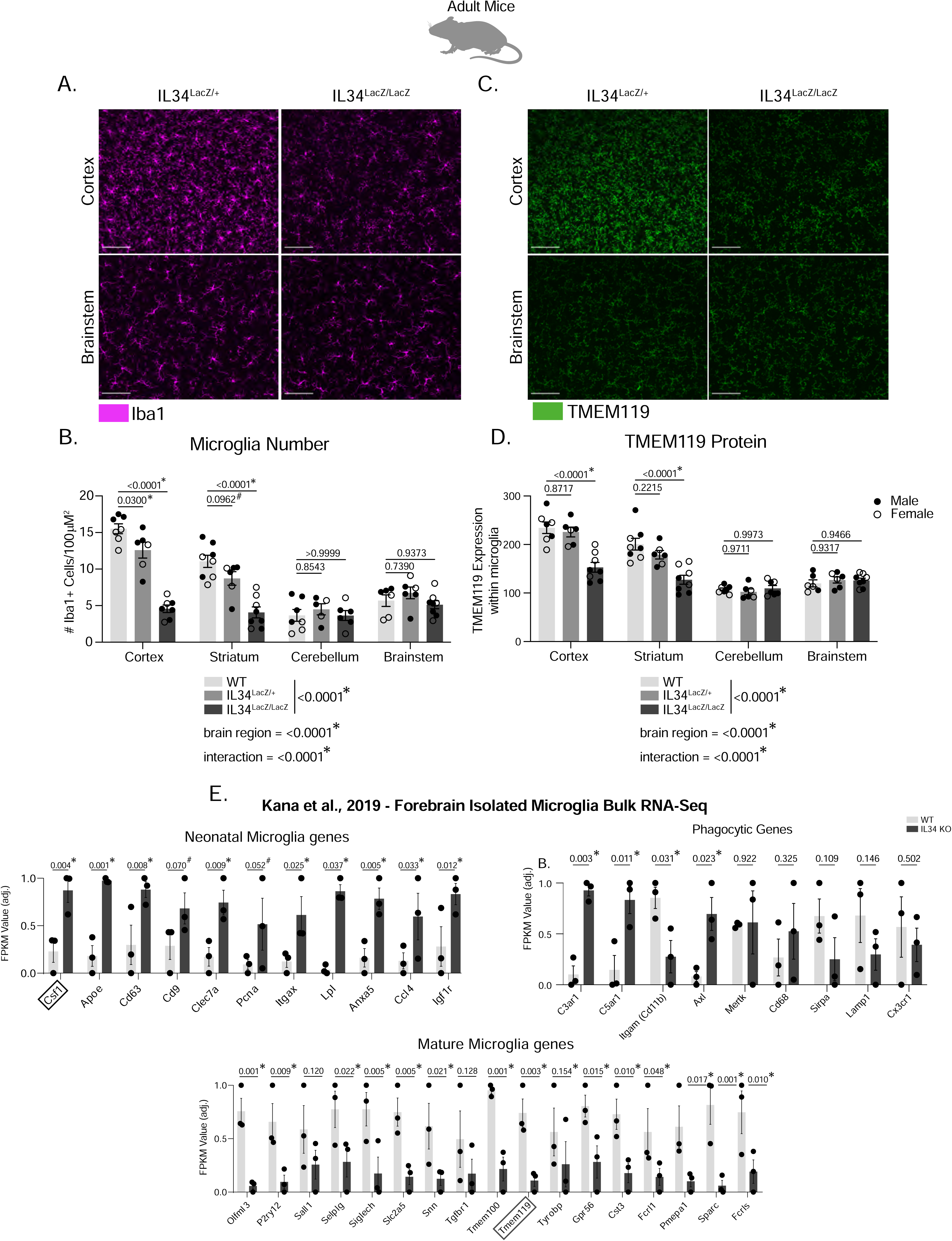
IL34 KO impacts microglia numbers and TMEM119 expression in adult cortex and striatum, but not cerebellum or brainstem. Related to Figure 2. (A and C) Representative images of Iba1 and TMEM119 stain in WT, IL34^LacZ/+^ and IL34^LacZ/LacZ^ mice from both the cortex and brainstem. Scale bar = 100µM. (B and D) Quantification of microglia number and TMEM119 mean gray value in the cortex, striatum, cerebellum, and brainstem from WT, IL34^LacZ/+^ and IL34^LacZ/LacZ^ mice. (n = 2-4 mice/sex/genotype, two-way ANOVA, Sidak’s post-hoc test, main effect of genotype and brain region and interaction in legend). (E) Scaled gene expression (FPKM) of embryonic/neonatal and adult microglia markers and candidate phagocytic genes from forebrain microglia from WT and IL34^LacZ/LacZ^ (IL34 KO) male mice. P values shown are from multiple t-tests function in Graphpad Prism. List of marker genes compiled from sequencing done in Bennett et al., 2016, Matcovitch-Natan 2014, and Li et al., 2019. Data from this figure was originally generated and published in Kana et al., 2019 and is accessible at GSE133362.

**Supplemental Figure 3.**
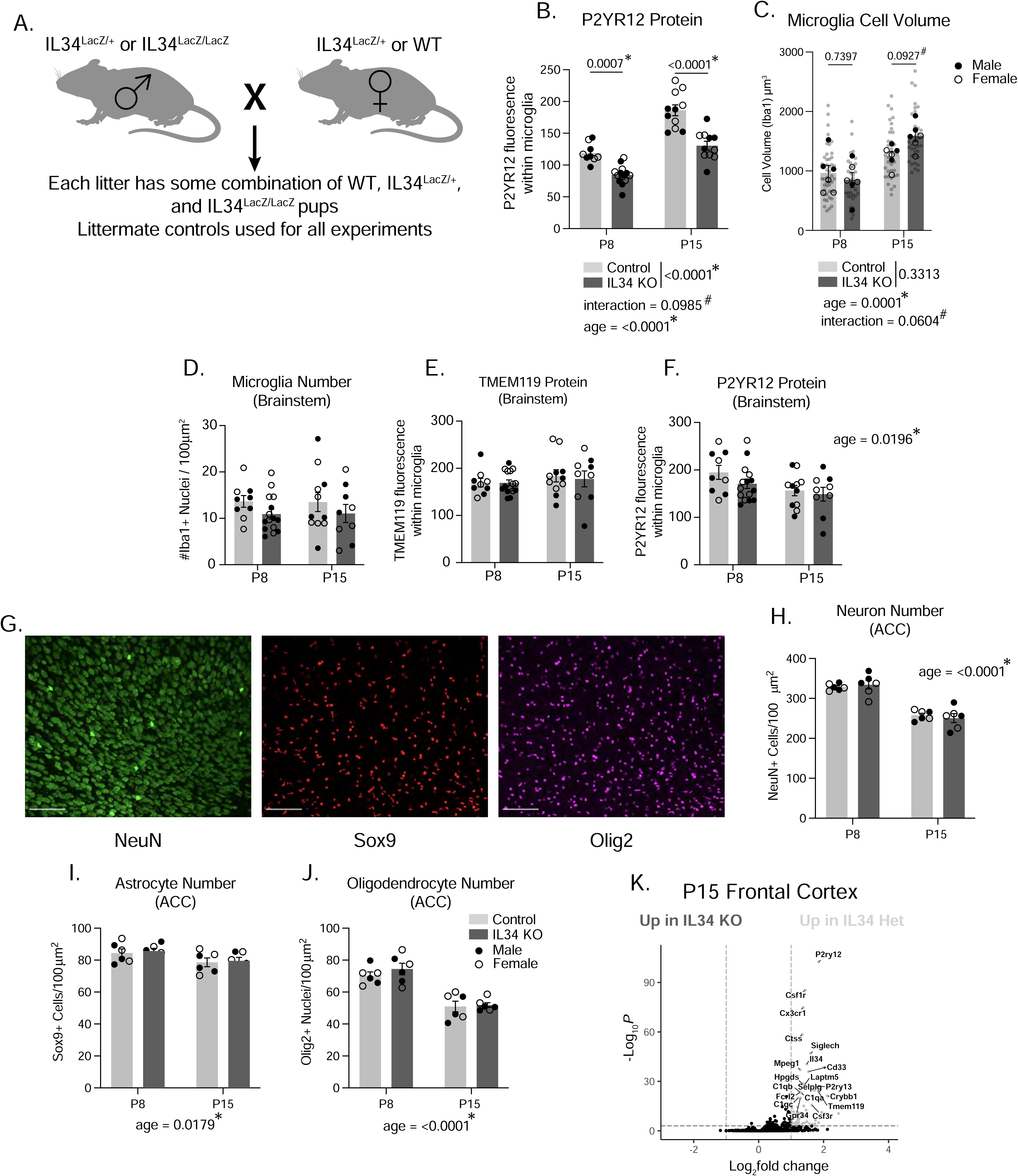
IL34 KO specifically impacts microglia in the forebrain. Related to Figure 2. (A) Representative schematic of IL34^LacZ/LacZ^ breeding scheme. (B) Quantification of P2YR12 mean gray value in the ACC of P8 and P15 IL34 KO and Control mice (n = 5-8 mice/sex/age/genotype, two-way ANOVA age x genotype, Sidak’s post-hoc test, main effect of genotype and age and interaction term in legend). (C) Quantification of microglia Iba1 cell volume from IMARIS 3D reconstructions. (n = 3 mice/sex/age/genotype, 4-6 cells analyzed per mouse, individual microglia represented by gray circles, animal averages represented by black dots, two-way ANOVA age x genotype, Sidak’s post-hoc test, main effect of genotype and age and interaction term in legend). (D-F) Quantification of microglia cell number, TMEM119 expression, and P2YR12 expression from the brainstem of P8 and P15 control and IL34 KO mice. (n = 5-8 mice/sex/age/genotype, two-way ANOVA age x genotype). (G) Representative images of NeuN, Sox9, Olig2 triple stain for quantification of neuron, astrocyte, and oligodendrocyte numbers/density. Scale bar = 100µM. (H-J) Quantification of Neuron, Astrocyte, and Oligodendrocyte numbers in ACC of P8 and P15 control and IL34 KO mice. (n=3 mice/sex/age/genotype, data shown are an average of 3 images taken from 3 sections of each mouse, two-way ANOVA age x genotype, main effect of age and interaction term in legend where significant). (K) Bulk RNASequencing was performed on a separate cohort of IL34^LacZ/+^ and IL34^LacZ/LacZ^ mice at postnatal day 15. (n=2 mice/sex/genotype, genes shown as significant passed a threshold of padj < 0.001 and LogFC > 1).

**Supplemental Figure 4.**
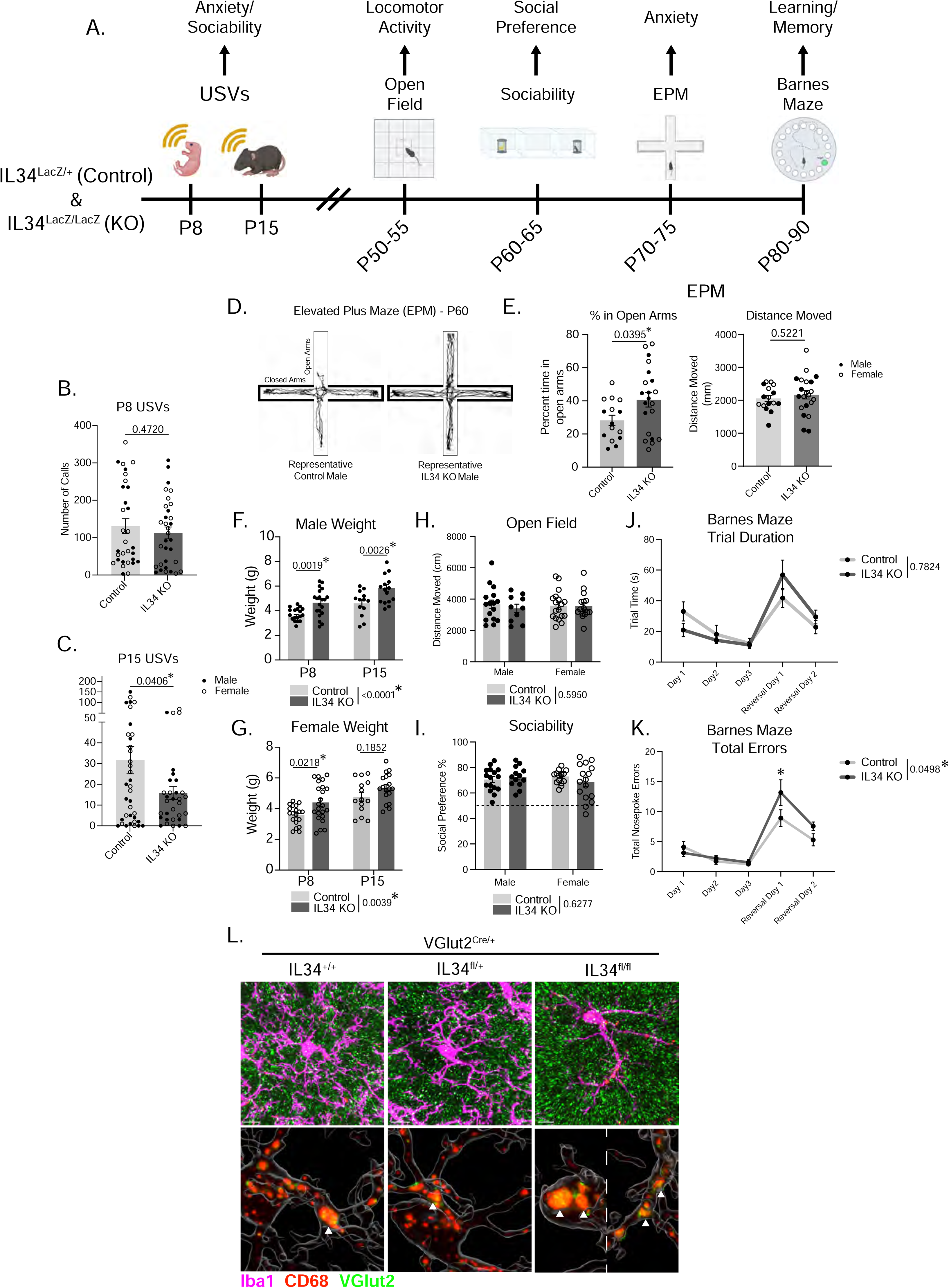
IL34 KO impacts pup weight, ultrasonic vocalizations, and anxiety in adulthood. Related to Figure 2. (A) Experimental timeline for IL34 KO behavioral phenotyping in development and adulthood. (B-C) Quantification of the total number of vocalizations made over the three-minute isolation period at P8 and P15 in IL34 KO and control mice. (n = 16-18 mice/sex/genotype, unpaired t tests). (D) Representative movement traces in the Elevated Plus Maze test in adult control and IL34 KO mice. (E) Quantification of percent time spent in open arms of the EPM and total distance moved over the 5-minute test (time spent in open arms / total test time * 100). (n = 6-12 mice/sex/genotype, unpaired t test). (F-G) Quantification of Male and Female pup weights from control and IL34 KO mice at P8 and P15. (n = 16-18 mice/sex/genotype, two-way ANOVA age x genotype, main effect of genotype in legend). (H) Quantification of total distance moved over a ten-minute period in the open field. (n = 10-16 mice/sex/genotype, two-way ANOVA sex x genotype, main effect of genotype in legend). (I) Quantification of social behavior from the three-chamber sociability assay. (n = 10-16 mice/sex/genotype, two-way ANOVA sex x genotype, main effect of genotype in legend). (J-K) Quantification of performance (trial time and errors) of male control and IL34 KO mice in the barnes maze across three training days and two reversal learning days. (n=8 male mice/genotype, two-way ANOVA day x genotype, main effect of genotype in legend, significance of errors on reversal day 1 from Sidak’s post-hoc test). (L) Raw IMARIS representative images of VGlut2 synaptic material engulfed in CD68+ lysosomes within microglia in the ACC of VGlut2^Cre^ IL34^+/+^, IL34^fl/+^, and IL34^fl/fl^ mice.

**Supplemental Figure 5.**
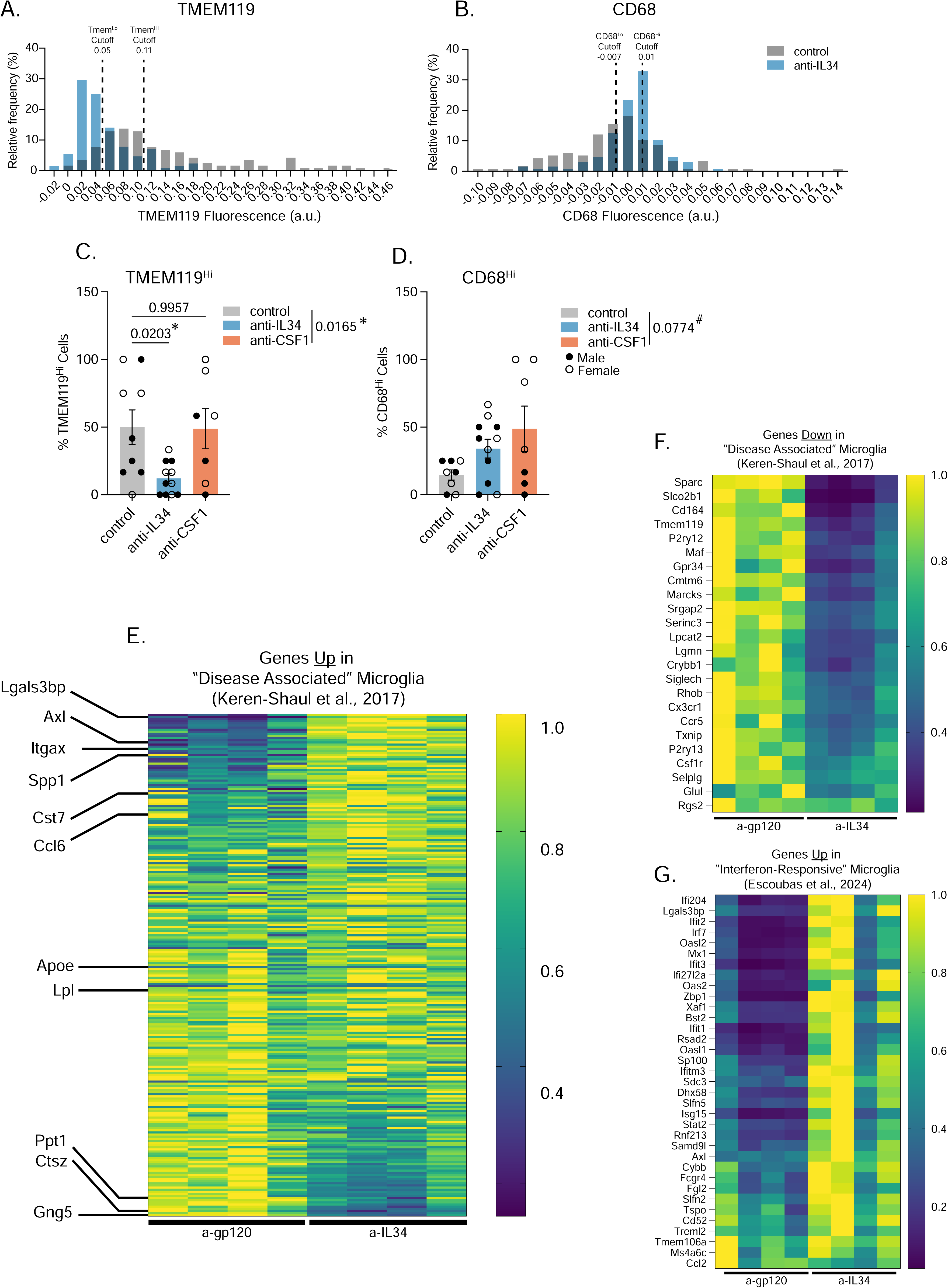
anti-IL34 microglia share some transcriptional similarities with disease-associated microglia and interferon-responsive microglia. Related to Figure 5. (A-B) Histograms of relative fluorescence of TMEM119 and CD68 stain within individual microglia. (C-D) Quantification of TMEM119^Hi^ and CD68^Hi^ microglia from control, anti-IL34, and anti-CSF1 mice. (n = 3-6 mice/sex/antibody, data shown are a percentage of 12 cells measured per animal across 3 images, one-way ANOVA, Sidak’s post-hoc test, main effect of antibody in legend). (E-G) Heatmaps of normalized TPM values between a-gp120 (control) and a-IL34 isolated microglia including all upregulated or downregulated genes from the “disease-associated” microglia cluster from Keren-Shaul et al., 2017, and all upregulated genes from the “interferon-responsive” microglia cluster from Escoubas et al., 2024.

**Supplemental Figure 6.**
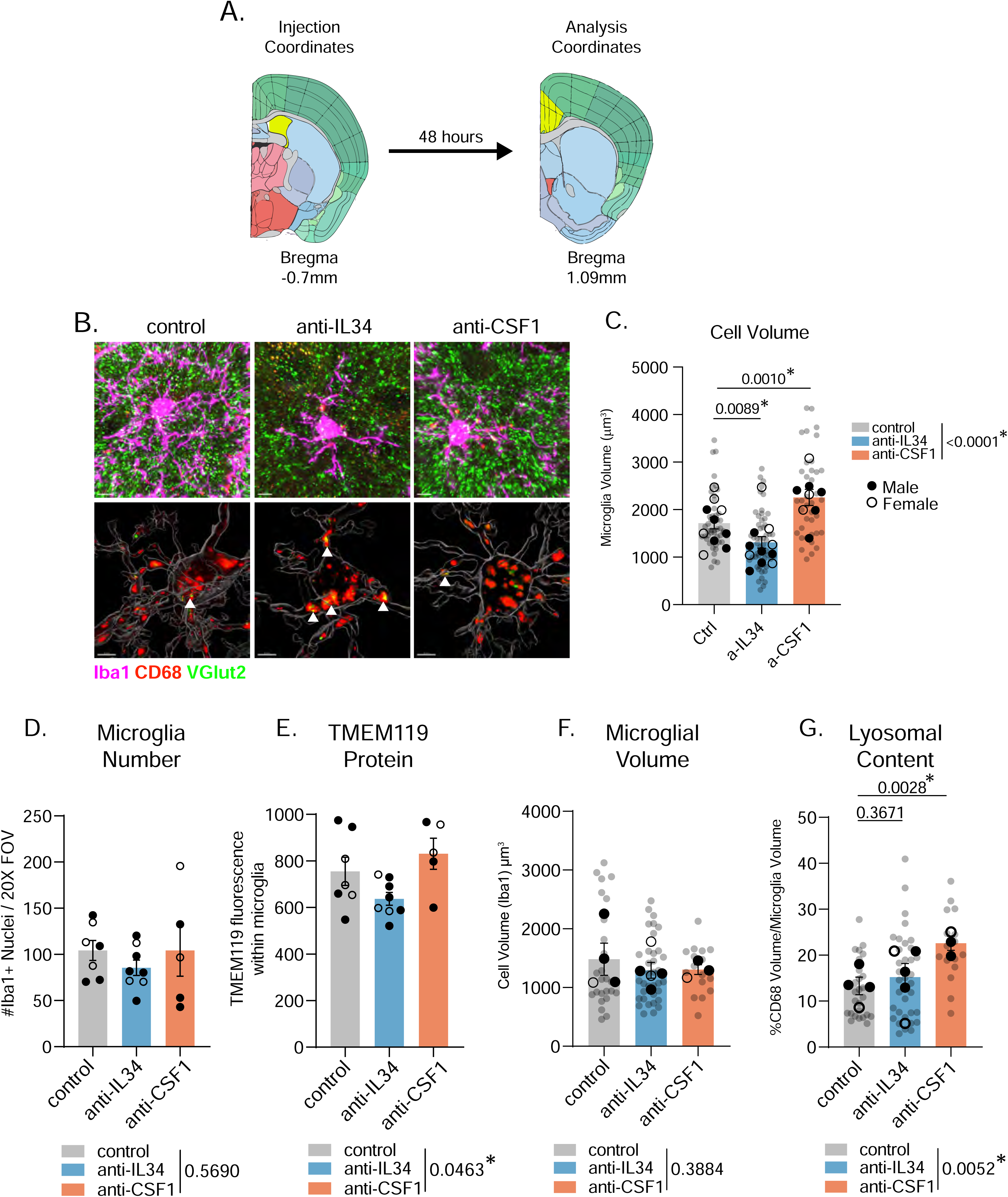
anti-IL34 does not impact cell number, but reduces TMEM119 expression and increases synaptic engulfment in the hippocampus. Related to Figure 6. (A) Representative atlas coordinates for lateral ventricle injection and ACC analysis. Images courtesy of Allen Brain Atlas. (B) Raw fluorescent representative images of microglia from control, anti-IL34, and anti-CSF1 mice. (C) Quantification of microglial cell volume. (n = 3-5 mice/sex/antibody, 4-6 cells analyzed per mouse, 141 cells total, individual microglia represented by gray circles, animal averages represented by black dots, nested one-way ANOVA, Sidak’s post-hoc test, main effect of antibody in legend). (D-E) Quantification of microglia number and TMEM119 protein in the hippocampus. (n = 2-5 mice/sex/antibody, one-way ANOVA, main effect of antibody in legend). (F-G) Quantification of microglia volume and lysosomal content in hippocampal microglia from control, anti-IL34, and anti-CSF1 mice. (n = 1-3 mice/sex/antibody, 4-6 cells analyzed per mouse, individual microglia represented by gray circles, animal averages represented by black dots, nested one-way ANOVA, Sidak’s post-hoc test, main effect of antibody in legend).

**Supplemental Figure 7.**
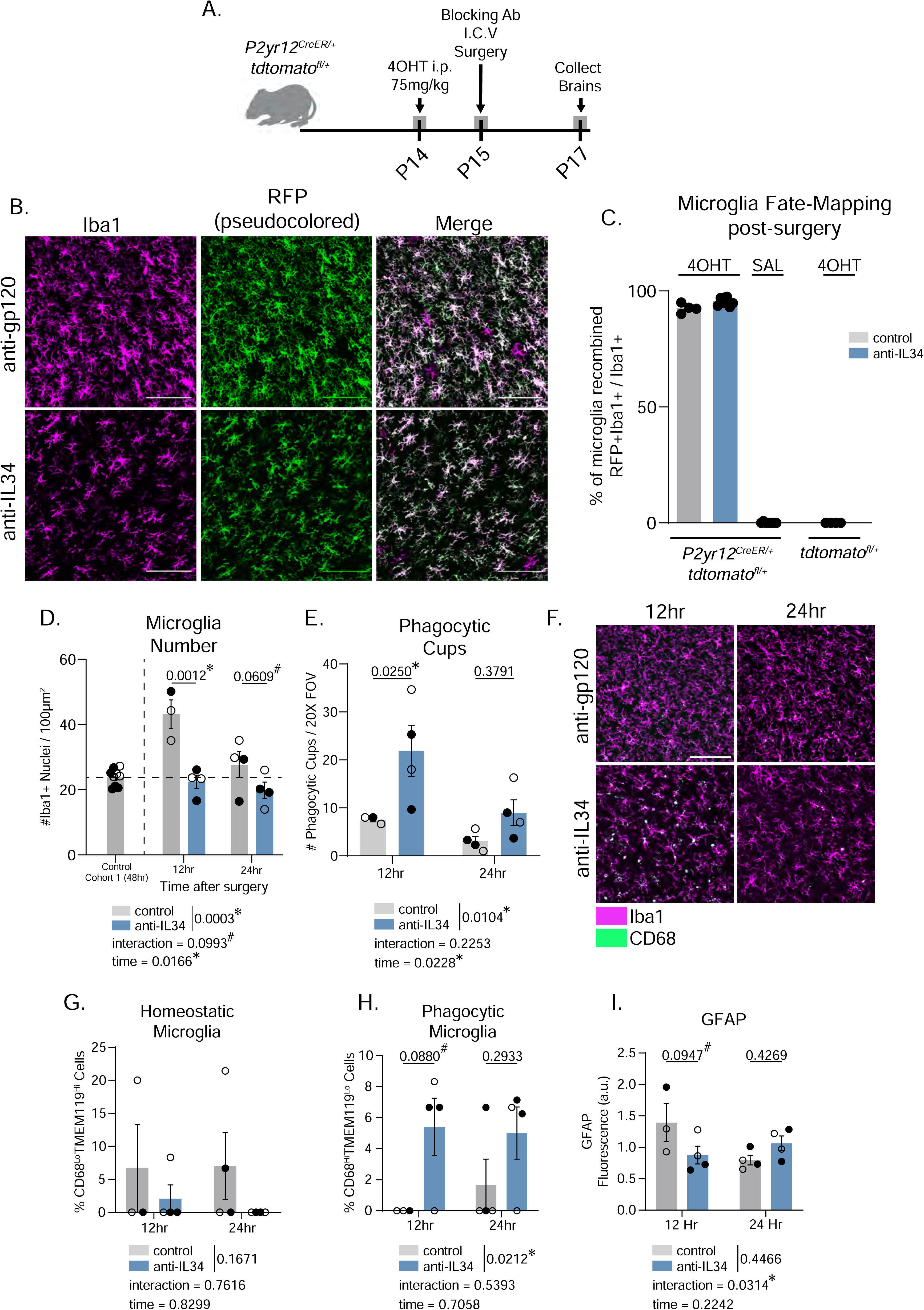
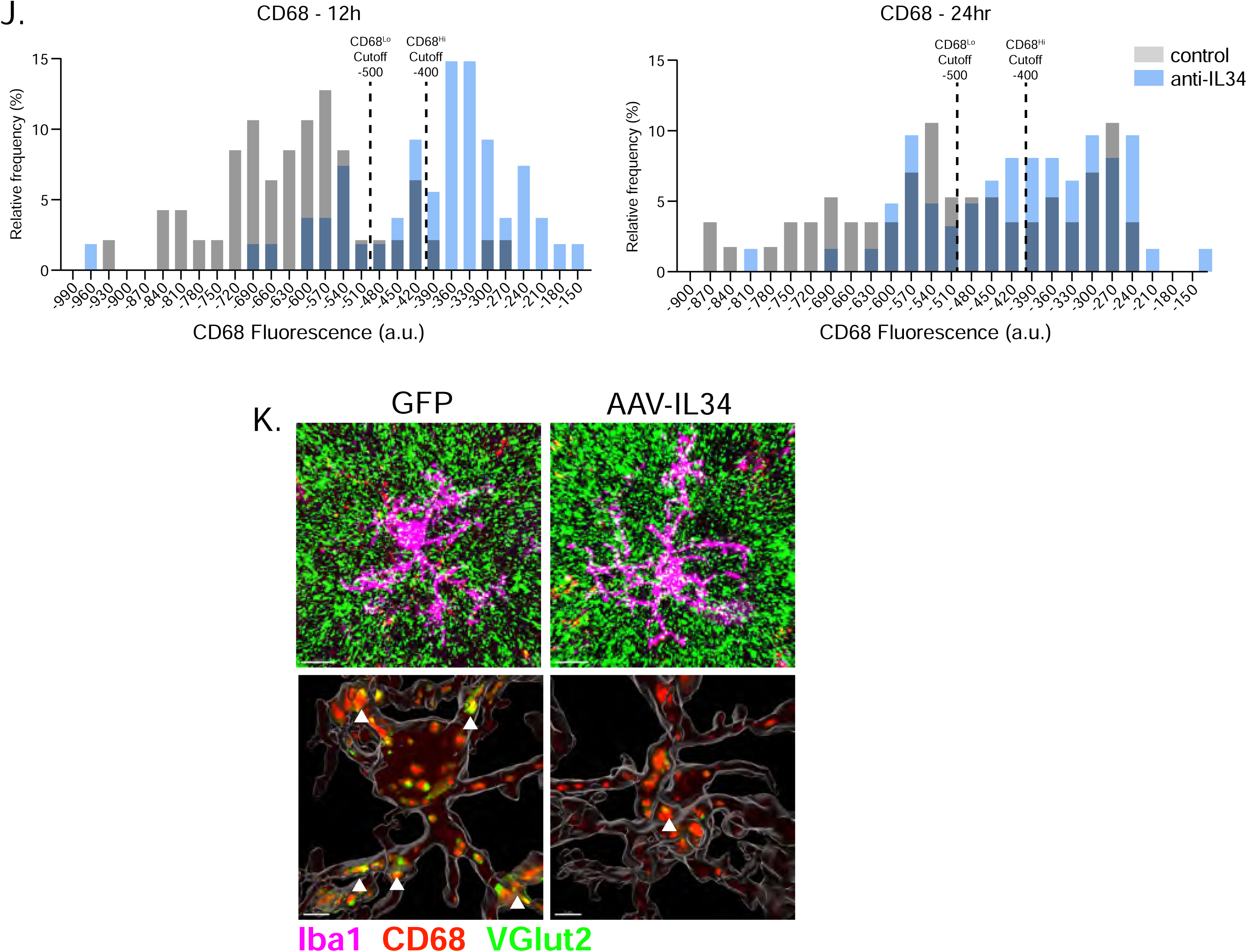
anti-IL34 treatment does not cause mass microglial death and repopulation. Related to Figure 4, 5, and 6. (A) Experimental schematic of 4-hydroxytamoxifen injection, blocking antibody surgery, and tissue collection. (B) Representative images of Iba1 and RFP staining in the ACC of mice treated with either the control or anti-IL34 blocking antibody. Scale = 100uM (C) Quantification of the percentage of microglia (total Iba1+ cells) that were also RFP+ following blocking antibody administration. (D) Quantification of microglia number in mice treated with a control or anti-IL34 blocking antibody 12- and 24-hours following surgery. Microglia number from controls from the original cohort collected 48 hours post-surgery is also shown. (n = 3-4 mice/timepoint/antibody, two-way ANOVA, Sidak’s post-hoc test, main effect of antibody and timepoint and interaction effect in legend). (E) Quantification of the number of phagocytic cups within 20X field-of-view. (n = 3-4 mice/timepoint/antibody, two-way ANOVA, Sidak’s post-hoc test, main effect of antibody and timepoint and interaction effect in legend). (F) Representative images of Iba1 and CD68 staining in mice treated with a control or anti-IL34 blocking antibody 12- and 24-hours following surgery. (G-H) Quantification of CD68^Lo^TMEM119^Hi^ homeostatic microglia and CD68^Hi^TMEM119^Lo^ phagocytic microglia in all four groups. (n = 3-4 mice/timepoint/antibody, data shown are a percentage of 12 cells measured per animal across 3 images, two-way ANOVA, Sidak’s post-hoc test, main effect of antibody and timepoint and interaction effect in legend). (I) Quantification of GFAP mean gray value in all four groups. (n = 3-4 mice/timepoint/antibody, two-way ANOVA, Sidak’s post-hoc test, main effect of antibody and timepoint and interaction effect in legend). (J) Histograms of relative fluorescence of CD68 stain within individual microglia at both timepoints. (K) Raw IMARIS representative images of VGlut2 synaptic material engulfed in CD68+ lysosomes within microglia in control GFP and AAV-IL34 mice.

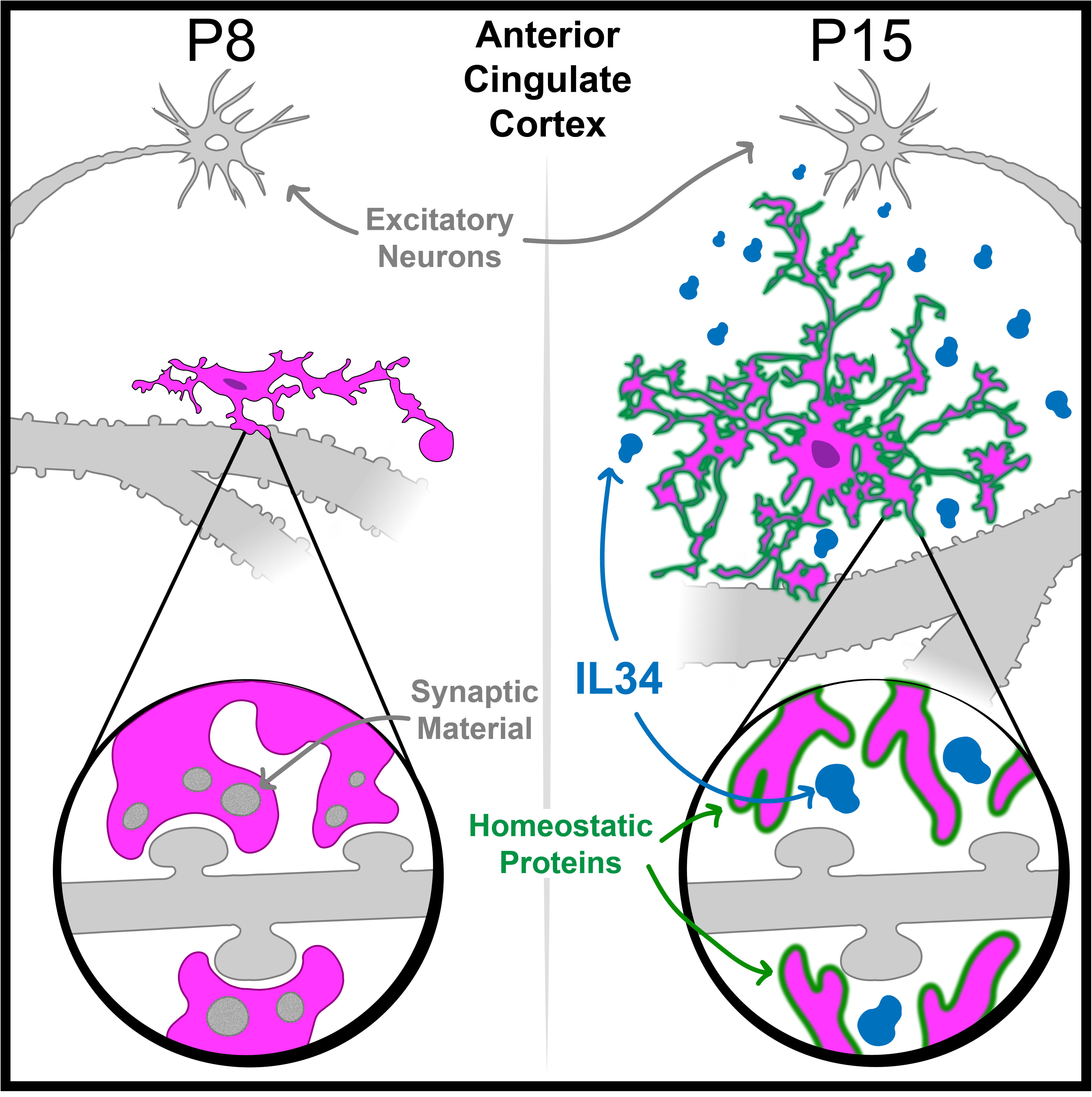

